# Lipid-mediated mechanism of drug extrusion by a heterodimeric ABC exporter

**DOI:** 10.1101/2025.02.26.640354

**Authors:** Qingyu Tang, Matt Sinclair, Paola Bisignano, Yunsen Zhang, Emad Tajkhorshid, Hassane S. Mchaourab

## Abstract

Multidrug transport by ATP binding cassette (ABC) exporters entails a mechanism to modulate drug affinity across the transport cycle. Here, we combine cryo-EM and molecular dynamics (MD) simulations to illuminate how lipid competition modulates substrate affinity to drive its translocation by ABC exporters. We determined cryo-EM structures of the ABC transporter BmrCD in drug-loaded inward-facing (IF) and outward-facing (OF) conformations in lipid nanodiscs to reveal the structural basis of alternating access, details of drug-transporter interactions, and the scale of drug movement between the two conformations. Remarkably, the structures uncovered lipid molecules bound in or near the transporter vestibule along with the drugs. MD trajectories from the IF structure show that these lipids stimulate drug disorder and translocation towards the vestibule apex. Similarly, bound lipids enter the OF vestibule and weaken drug-transporter interactions facilitating drug release. Our results complete a near-atomic model of BmrCD’s conformational cycle and advance a general mechanism of lipid-driven drug transport by ABC exporters.

## INTRODUCTION

Ubiquitous across all kingdoms of life, ABC transporters traffic molecules across cell membranes in both directions by directly consuming ATP (*1–3*). A subclass of these transporters, ABC exporters, specializes in exporting structurally and chemically diverse substrates, often cytotoxic molecules, thereby imparting a multidrug resistance phenotype (*3–5*). The canonical alternating access mechanism of the ABC exporter class invokes the population of a suite of conformations powered by ATP binding and hydrolysis in the dedicated nucleotide-binding domains (NBDs) (*6–8*). Through the underlying conformational changes, substrate molecules bind an inward-facing (IF) vestibule cradled by the transmembrane domains (TMDs) and are then extruded following a switch of the vestibule orientation to face the extracellular side. For multidrug ABC exporters, amphipathic substrates can bind from either the inner leaflet of the bilayer or the cytoplasm and are released to the outer leaflet of the bilayer or the extracellular milieu (*9, 10*).

Insight into the determinants of polyspecific binding has been gleaned from multiple drug-bound IF conformations of multidrug ABC transporters (*11, 12*). However, there is a paucity of drug-bound outward-facing (OF) structures (*7, 10, 13, 14*), although a low-resolution OF structure has been reported for substrate-bound BmrA(*15*). In addition, P-glycoprotein (ABCB1) was capture in IF and purportedly OF structures bound to a synthetic substrate that was trapped via disulfide bonds in cysteine mutant backgrounds. The covalent trapping, the unexpected location of the synthetic substrate and the occluded nature of the conformation raise questions regarding its mechanistic identity(*16*). Moreover, neither Pgp structures were determined in a wild-type background and in a lipid bilayer environment(*15, 16*). The dogma in the field associates the release of substrates by ABC transporters to a reduction in affinity in the OF conformation. However, addressing the unanswered central question of how interactions with the transporter are weakened enough to drive release following the power stroke requires high resolution structures of drug-bound transporters in a native-like environment.

Here we illuminate a lipid-driven mechanism of substrate release through an integrated single particle cryo-EM and MD simulations investigation of the heterodimeric multidrug ABC exporter BmrCD from *B. subtilis*. BmrCD has been the subject of extensive functional, biochemical, spectroscopic and cryo-EM investigations in lipid bilayers yielding a profound understanding of multiple facets of its mechanism (*8, 17–20*). The work presented in this paper addresses the missing critical element of how drugs are released to the extracellular side and proposes a general model of multidrug export.

## RESULTS

### Cryo-EM structure of Hoechst-bound BmrCD in an OF conformation

We have previously reported a series of BmrCD conformations in lipid nanodiscs including multiple ATP- and drug-bound IF intermediates and an occluded (OC) conformation (*8*). Driven by the evidence from double electron-electron resonance (DEER) spectroscopy that an OF conformation is populated (*8, 17*), we identified 2D images that profile such a structure featuring an open extracellular side (Fig. S1). Subsequent 3D classification, based on OC particles, from a dataset of the WT transporter in an ADP-Vi trapped biochemical preparation yielded a small subset of OF particles (Fig. S2). Biochemical, spectroscopic and structural data established that under vanadate trapping following ATP hydrolysis, ABC exporters populate a high-energy, post-hydrolysis state, often in a predominantly OF conformation (*8, 17, 21*). A final map with a resolution of 3.1Å of an OF conformation was obtained (Fig. 1A and Fig. S2) following non-uniform and local refinement. The OF model (Table S1, and Figs. S2 and S3) features a V-shaped vestibule open to the extracellular side and the outer leaflet of the bilayer. In this OF vestibule, two Hoechst molecules pack in an antiparallel conformation (Figs. 1A and 2). In addition to either 2 molecules of ATP or an ATP/ADP-Vi in the NBDs, twenty lipid molecules are observed to decorate the intracellular and extracellular halves of the transporter. Lipids were conspicuously bound to each side of the V-shaped interface of BmrC and BmrD at the extracellular side (Fig. 1A, and Fig. S2G). Previous DEER analysis (*17*) demonstrated that ATP hydrolysis is required for population of the OF conformation, thus, we have modeled the consensus site as bound to ADP-Vi in the cryo-EM structure.

**Fig. 1.**
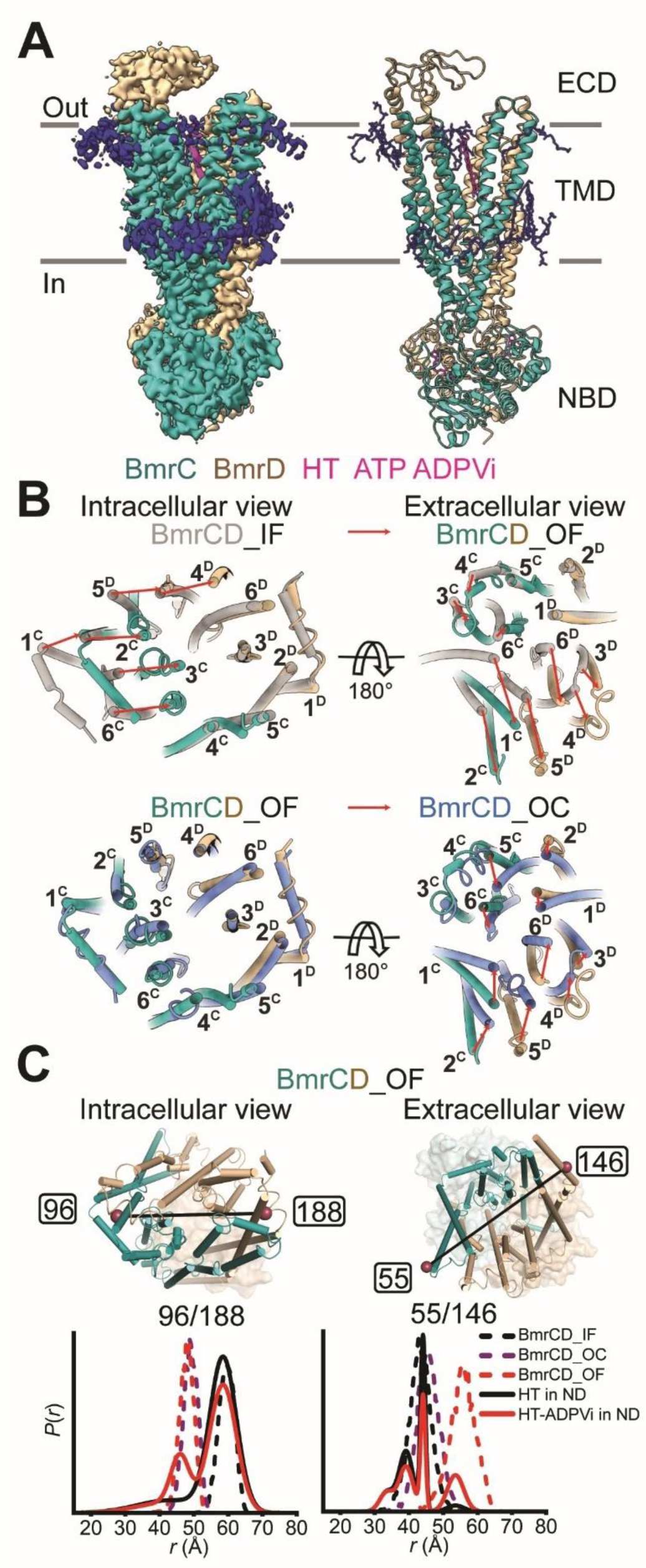
Structure of Hoechst-bound BmrCD in an outward-facing conformation. (**A**) Cryo-EM map and structure model of BmrCD OF conformation bound to Hoechst. The extracellular domain (ECD), transmembrane domain (TMD), and nucleotide-binding domain (NBD) are indicated. BmrC and BmrD are shown in light sea green and tan. Hoechst (HT) and ATP/ADPVi are shown as sticks in magenta, and lipids are blue, hereafter if not indicated. (**B**) Conformational changes of the TMD intracellular (left panels) and extracellular sides (right panels) between BmrCD_OF, BmrCD_IF (PDB 8FMV), and BmrCD_OC (PDB 8T1P). All alignments are based on BmrD. BmrCD_IF and BmrCD_OC are shown in grey and slate, respectively. Large movements are indicated by red arrows. (**C**) DEER distance distributions for spin-label pairs on the intracellular (96^BmrC^/188^BmrD^) and extracellular (55^BmrC^/146^BmrD^) sides in BmrCD. Upper panels: close-up views of BmrCD_OF highlighting spin label pairs. TMD helices are shown as cylinders, ECD and NBD are shown as transparent surfaces. The spin-labeled locations are shown by raspberry spheres. Lower panels: experimental distance distributions are shown in solid lines. Simulate distance distributions derived from cryo-EM structures by MDDS are shown in dashed lines.

**Fig. 2.**
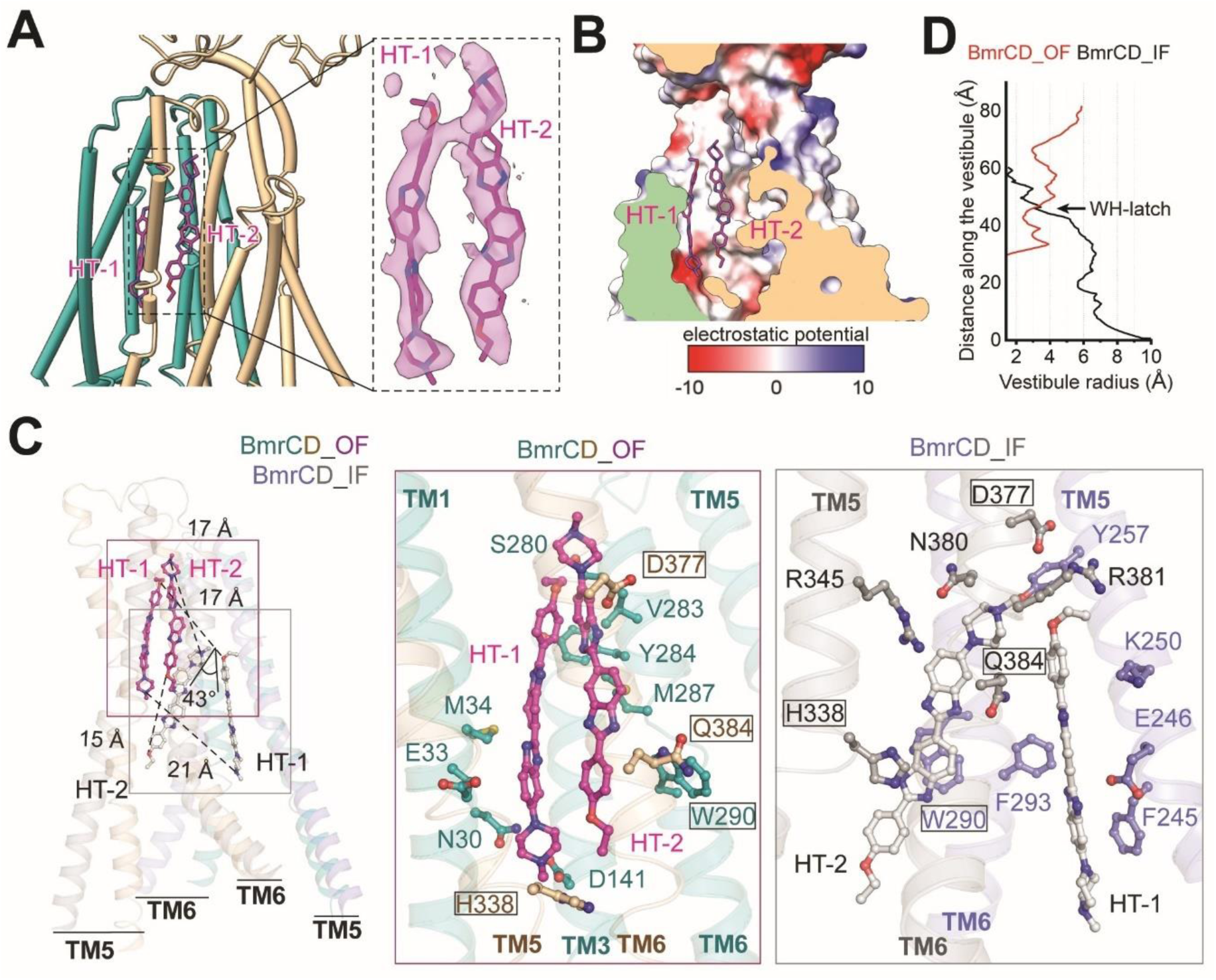
Two Hoechsts are bound to the OF structure. (**A**) Close-up view of the two bound Hoechsts represented by magenta sticks and colored by heteroatom. The cryo-EM density of Hoechsts are shown in the inset. (**B**) Electrostatic potential surface of the Hoechsts-binding chamber of BmrCD_OF. (**C**) Perspective on Hoechst movement from the IF to the OF conformation. BmrC and BmrD of IF are shown in slate and grey and Hoechsts of IF are represented by grey sticks colored by heteroatom. Left panel: BmrCD OF and IF are superimposed based on BmrD. The shift in distances and angles of the Hoechsts are noted. Transmembrane helices TM5 and TM6 from both BmrC and BmrD are shown in transparent cartoon. Middle and right panels: key residues that stabilize Hoechsts in the OF and IF conformations are labeled, residues in common are boxed. (**D**) Radius measurement of the vestibule in the OF and IF structures reveals that WH-latch (W290^BmrC^H338^BmrD^-latch)(*8*) is located at the bottleneck of the chamber.

Superposition of Hoechst-bound BmrCD OF (BmrCD_OF) and IF (PDB 8FMV hereafter referred to as BmrCD_IF) (*8*) conformations reveals that the closing of the intracellular side is mainly contributed by BmrC rearrangement, whereas opening the extracellular side is executed by BmrD reconfiguration. Specifically, TM1, TM2, TM3, and TM6 of BmrC, along with TM4 and TM5 of BmrD, exhibit substantial displacement, resulting in the closure of the intracellular vestibule, with distance changes ranging from 8 to 12 Å. Concomitantly, the extracellular side of the TM helices move outward, facilitating the formation of an OF vestibule. Opening of the extracellular side is primarily driven by TM1 and TM2 of BmrC and TM3, TM4, TM5, and TM6 of BmrD with distance changes ranging from 6 to 15 Å (Fig. 1B, upper panels). These distance changes are validated by DEER distance measurements in the corresponding lipid nanodiscs (*8, 17, 21*). A minor population at the larger distance of the extracellular pair 55/146, observed in the vanadate-trapped post-hydrolysis state(*8*) (Fig. 1C, solid red line), overlaps the predicted distance distribution from the OF structure (Fig. 1C, dashed red line) further reinforcing the mechanistic identity of this structure. Superposition of BmrCD_OF and OC (PDB 8T1P hereafter referred to as BmrCD_OC) (*8*) conformations reveals that the closing of the extracellular side at this step is mainly contributed by BmrD rearrangement (Fig. 1B, lower panels). In addition, the asymmetry of TMD between BmrC and BmrD is reduced in the OF compared to the IF and OC structures (Fig. S4).

The conformational changes propagate to the extracellular domain (ECD) that blocks the TMD in the IF conformation. Four loops, residues 92-96, 103-106, 111-116, and 120-124, are displaced approximately 4-5 Å in the IF to OF transition, while they move back 4-7 Å in the transition from OF to OC (Fig. S5A). These conformational changes indicate that ECD movement is coupled to Hoechst extrusion. The local structural changes of the ECD have a profound effect on the electrostatic surface which could play a role in funneling the Hoechst molecules out of the transporter. Specifically, the positive central patch (black circle) of the ECD in OF can repel the positively charged Hoechst, whereas the two negative side patches (blue and yellow circles) attract it, which may guide the Hoechst extrusion. Notably, these charge distributions are reversed in the IF and OC conformations (Fig. S5B). Similar to the TMD, distance distributions for spin label pairs in the ECD, predicted from the OF structure, superimposes on the experimental data further confirming the conformational changes at these sites (*8*) (Fig. S5C).

### Interactions of Hoechst with the outward-facing vestibule of BmrCD

Clear density of two Hoechst substrates (labeled HT-1 and HT-2) enables a detailed analysis of their interactions with the transporter in the OF conformation (Fig. 2, A and B). The two Hoechst molecules are located between TM1, TM3, and TM6 of BmrC and TM5 and TM6 of BmrD. Specifically, D141 of BmrC and D377 of BmrD hydrogen-bond with the piperazine group of HT-1 and the benzimidazole group of HT-2, respectively. The pivotal role of D141 is consistent with previous results showing that its alanine substitution abrogates Hoechst stimulation of ATP hydrolysis in BmrCD (*21*). Y284 of BmrC interacts with HT-1 via a π-π interaction, and V283 of BmrC forms hydrophobic interaction with HT-2. M34 and M287 of BmrC hydrophobically interact with the two Hoechsts separately (Fig. 2C, middle panel). Similar to the IF conformation, the major residues contributing to Hoechst binding are from BmrC, suggesting that BmrC interactions guide Hoechst during transport (Fig. 2C). W290 and H338 of BmrC, D377 and Q384 of BmrD appear to interact with Hoechst in both conformations (Fig. 2C).

Comparison of the IF and OF conformations (Fig. 2C) captures the translocation of the two Hoechst molecules by around 15 Å to 21 Å from the lower part to the upper part of the TMD. In the IF conformation, the two Hoechsts form an angle of 43°, while in the OF conformation, they become almost antiparallel. The alternating access of the vestibule is contributed by the intracellular and the extracellular sides of BmrD TM5 and BmrC TM6 which move to close the entrance and open the exit simultaneously. The substrate binding vestibule’s volume is reduced from 7882 Å^3^ to 4995 Å^3^ during the IF to OF transition. The fulcrum of this transition (Fig. 2D) appears to be located in the vicinity of a previously(*8*) identified ionic latch between W290^BmrC^ and H338^BmrD^, both critical residues for binding of Hoechsts (Fig. 2C, middle and right panels). Interestingly, inspection of the putative OF conformation of mPgp(*16*) shows that TM6 TM11 (equivalent to TM5 in the second half of Pgp) contribute to substrate binding in both OF and IF conformations with the equivalent residue to W290 (F339) in mPgp contacting the substrate (Fig. S6). This residue is also conserved in hPgp, and the bacterial heterodimers TmrAB and TM287/288 (*15, 22*) (Fig. S6).

### MD simulations uncover the pivotal role of lipids in Hoechst release

We simulated the dynamics of the lipid molecules visualized in the cryo-EM structure with particular focus on the lipid molecule, labeled POV15, which is poised to enter the extracellular cleft of the protein. Our simulations show a rapid descent of this lipid into the cleft. Its entrance is captured in kernel density estimation analysis which shows regions of high occupancy or density, revealing locations accessible to membrane phospholipids (Fig. 3, A and B). In one simulation replica the lipid-binding event is aborted, while in the other 3 replicas it continues until the lipid has fully entered the transporter OF cavity.

**Fig. 3.**
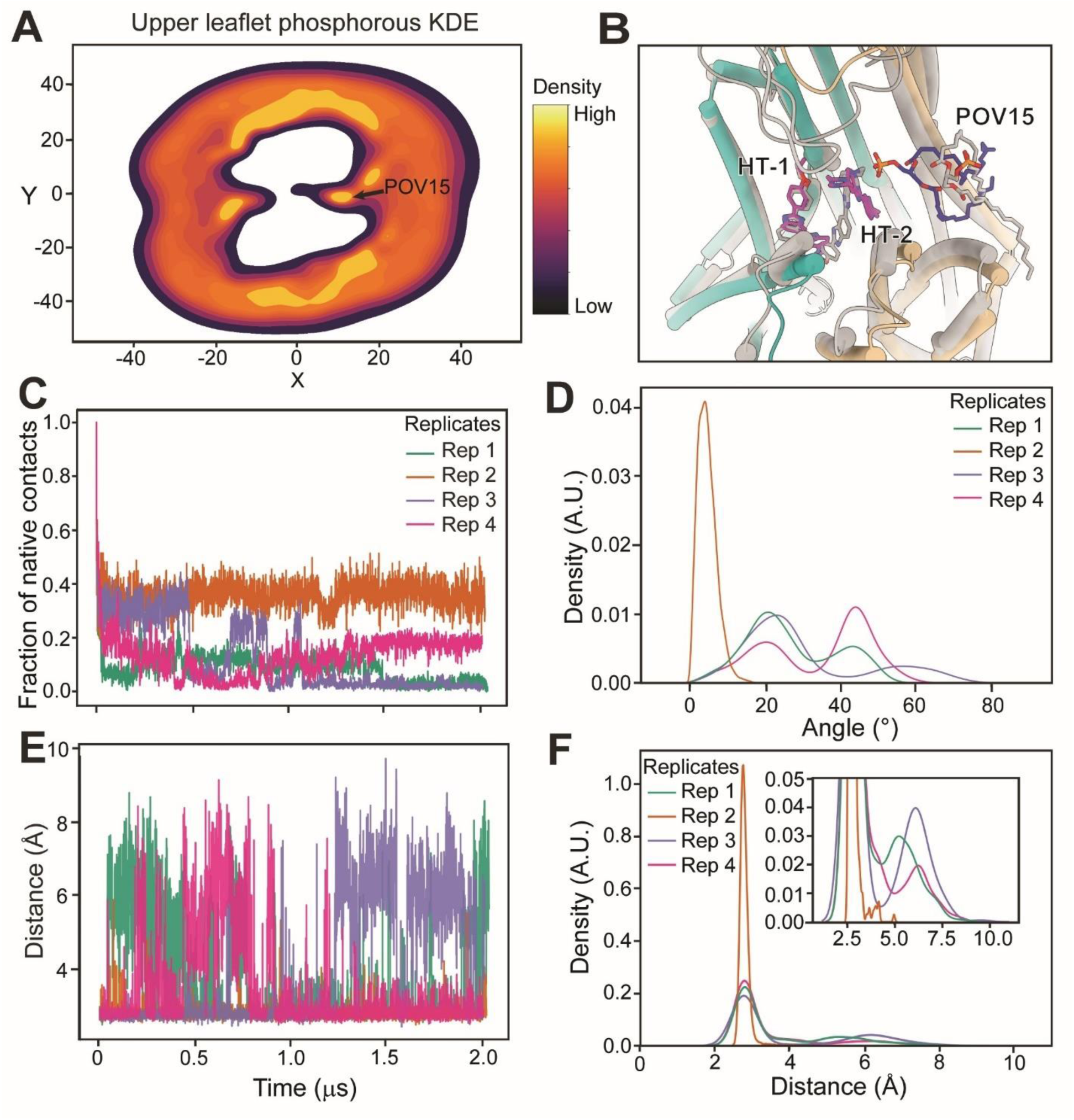
MD simulations of lipids bound to BmrCD_OF. (**A**) Kernel Density Estimation analysis. Lipid phosphorous atoms in the outer leaflet have been binned in their XY projections with high density appearing yellow and low density appearing dark purple. (**B**) Molecular rendering of POV15 entering into the cleft and attracting Hoechst molecules. Grey and color models are the MD simulation results at 0 ns and 140 ns, respectively. (**C**) Time series of fraction of native contacts. Each color in the trace belongs to an independent replica simulation. Native contacts are defined as any protein residue with heavy atoms within 4.5 Å of the Hoechst molecules in the initial structure. (**D**) Distribution of Hoechsts tilt angles as measured with respect to the positive Z-axis. This was measured by defining the vector going from the piperazine to the ethoxy group of Hoechst and measuring the angle formed with the vector (0, 0, 1). (**E**) Time series for the salt-bridge distance between E33 and R345 of BmrC and BmrD, respectively. Each trace represents an independent replica. (**F**) Distribution of E33^BmrC^-R345^BmrD^ salt-bridge distances over all the simulation.

One consequence of the lipid entrance into the vestibule is the perturbation of the bound Hoechst molecules, highlighted in native contact analysis (Fig. 3C). While in the cryo-EM density the Hoechst molecules adopt roughly vertical, antiparallel conformations, the presence of the negatively charged lipid induces a strong tilt for both HT molecules (Fig. 3D). This tilting motion of the drug molecules breaks several of the stabilizing interactions formed between the two chains of BmrCD and Hoechst molecules. One such critical residue in substrate binding, D141 of BmrC(*21*), is also implicated in the observed lipid-mediated substrate dislodging. As the lipid weakens protein substrate contacts, D141 continues to hold tightly onto the charged end of HT-1, heavily contributing to the tilting behavior observed as lipids tug on the opposite end of HT-2. As POV15 begins pulling on HT-2, we also observe a change in the electrostatic interactions between BmrCD and Hoechst (Fig. S7A). This energetic perturbation occurs one substrate at a time and correlates with structural rearrangement of TM1 of BmrD (Fig. S7, B and C).

In another simulation replica we observed HT-1 beginning to depart the original binding site into the membrane, opposite of POV15 (Fig. S7, D and E). This lipid-substrate interaction helps to break the D141-HT-1 salt bridge as well as the nearby E33^BmrC^-R345^BmrD^ salt bridge, allowing E33 to now form a new salt bridge with the exposed charged nitrogen of HT-1 (Figs. 3, E, F, and S8A). Concurrently, HT-2 terminal piperazine group forms a salt bridge to E51 of BmrD, causing the substrate to rapidly translocate up towards the ECD domain (Fig. S8B). The piperazine group then is passed off to the nearby E75 of the ECD, while E51 goes on to form a salt bridge with the benzimidazole ring in the Hoechst molecule.

While HT-2 is traveling up towards the ECD domain, the ethoxy group of the molecule glides out of the extracellular cleft, an event accompanied with a correlated opening of the ECD domain (Fig. S8, C and D). In summary, we observe an unbinding motion which is initiated by the entry of POV15 into the transporter chamber. Shortly after lipid entry, HT-2 is engaged in a pattern of electrostatic and hydrophobic interactions with the lipid, allowing HT-1 to dislodge and presumably diffuse into the nearby membrane. This motion provides a potential mechanism for the appended ECD domain in substrate unbinding and may explain why BmrCD contains this structural domain relative to other type IV ABC transporters.

### MD simulations reinforce the pivotal of D141A in substrate release

In light of the previous experimental data and the above simulations of Hoechst dynamics in the OF conformation, we further probed the role of D141 in the release mechanism by MD simulations in a background where this residue is substituted by alanine. We tracked the distance between the center of mass (COM) of HT-1 or HT-2 and COM of residue 141 over WT and D141A simulations (2 µs). As shown in Table S2 and Fig. S9, A and B, HT-1 maintained a distance of 4.0 ± 1.7 Å to D141, in the WT with transient fluctuations generally under 2.0 Å (n = 3 replicates).

In contrast, in the D141A mutant, the distance to A141 drifted to a 7.5 ± 2.5 Å, occasionally even reaching above 14 Å. While less considerable, a similar pattern arose for HT-2: compared to a distance of 8.4 ± 2.7 Å in the WT, we observed a distance of 12.0 ± 1.8 Å for D141A. The RMSD of each ligand relative to its initial bound conformation (Fig. S9, C-F) further substantiates the impact of D141 removal. In WT, the average RMSD of HT-1 plateaued to 3.8 ± 1.0 Å (n = 3 replicates), while the mutant showed larger fluctuations, reaching up to 12.3 Å, with an average RMSD of 6.1 ± 3.1 Å (n = 3 replicates). Likewise, HT-2 in WT settled at an RMSD of 3.9± 1.6 Å, whereas in the D141A mutant it reached 4.6 ± 1.3 Å during the same period. These data demonstrate that, once D141 is removed, both HT-1 and HT-2 lose a key electrostatic anchor and adopt more mobile binding poses that deviate from the original binding pose more wildly. Consistent with these shifts, ionic bond analysis (Fig. S9, G and H) shows that D141 in the WT established recurring salt-bridge interactions with HT-1, but after mutation, these contacts disappeared, prompting the ligands to form compensatory electrostatic interactions with other negatively charged residues (e.g., E33 in BmrC).

Overall, analysis of the distances, RMSD and ionic interactions, strongly indicate that D141 is a pivotal anchor for HT-1 and HT-2, and when it is replaced by a non-charged residue (D141A), the ligands experience increased fluctuations and less stability in the binding pocket. These collectively drive HT-1 and HT-2 to shift away from the canonical binding region. While we did not observe full unbinding of the ligands from their binding pocket over the simulation timescales, the observed increased fluctuations after introducing the mutation clearly reflect an increased unbinding probability.

### Lipid binding to the IF conformation

The discovery of the role of lipids in the release of Hoechst from the OF conformation prompted us to investigate if lipids play a role in Hoechst binding to the IF conformation. Searching the same ADP/Vi-trapped dataset identified two IF conformations at a resolution of 3.4 Å, both with lipids bound in proximity to the two Hoechst molecules (Figs. 4A, S10, S11, and S12, Table S1). In one IF conformation, hereafter referred to as BmrCD_IF1, a PA molecule, (labeled as POV4), directly interacts with H184 of BmrC (Fig. 4, B and C, middle panels) which is part of an ionic “latch” that involve its interaction with R304 side chain, as we previously noted(*8*). The latch closes upon the binding of the two Hoechst molecules manifested by the distance between the two side chains(*8*). In BmrCD_IF1, the latch is in the open state as the lipid engages H184. In the second identified IF conformation, BmrCD_IF2, a lipid molecule (POV23) is bound in the substrate binding vestibule with the latch in the closed position. In addition, another lipid (POV4) is poised nearby (Fig. 4, B and C).

**Fig. 4.**
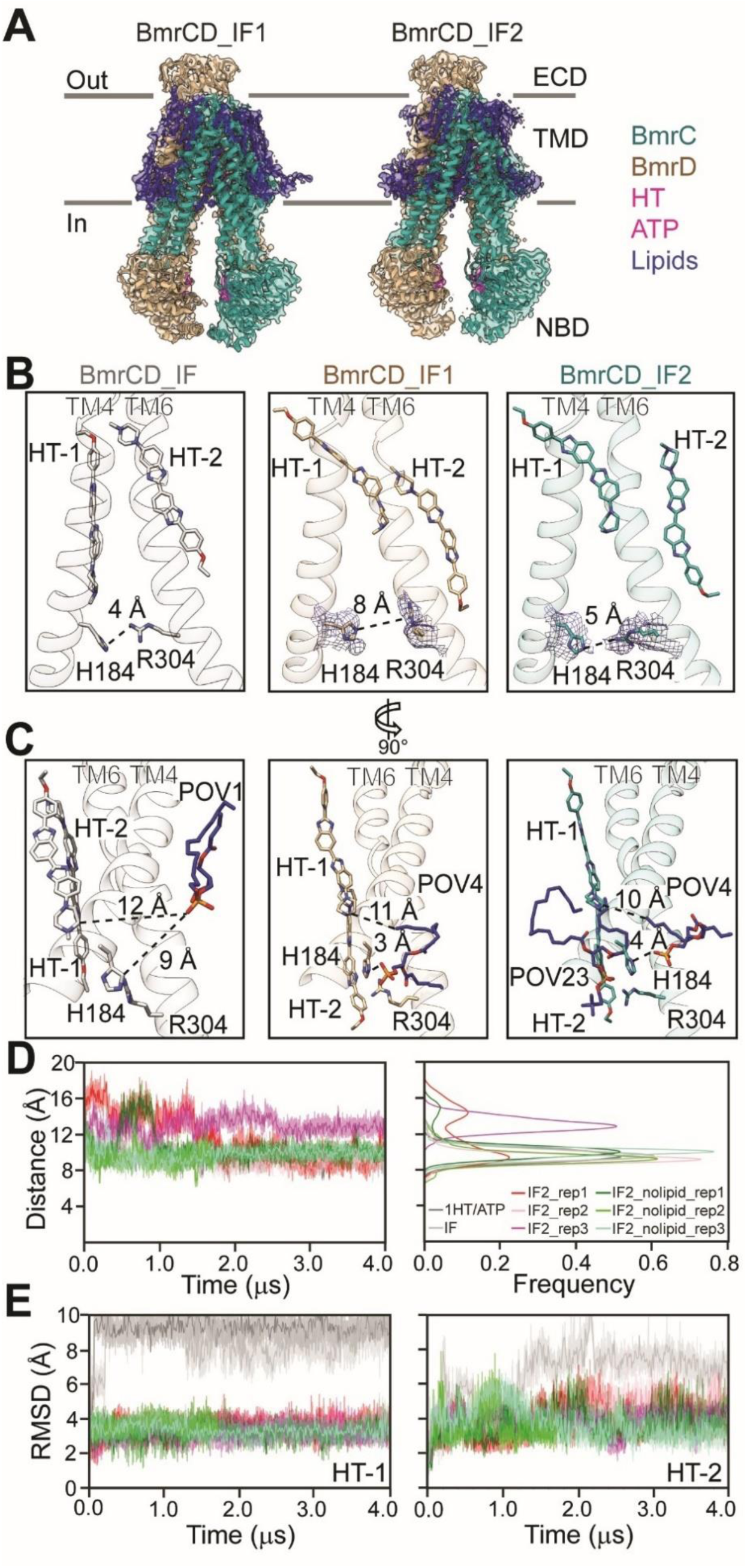
RH latch controls lipid entry the substrate-binding chamber. (**A**) The structure models and density maps of BmrCD_IF1 and BmrCD_IF2. Density maps are shown as transparent surface and color scheme is the same as Fig. 1A. (**B**) The RH-latch (R304^BmrC^-H184^BmrC^ latch) densities for BmrCD_IF1 (middle panel, tan), and BmrCD_IF2 (right panel, light sea green) are shown with side-chain distance indicated. BmrCD_IF (PDB 8FMV, left panel, grey) is shown for comparison. (**C**) The lipid molecules close to the RH-latch are shown in blue sticks colored by heteroatom with the distance highlighted. The lipids code is from the corresponding structure. (**D**) RH-latch monitored by MD simulations. Time series (left panel) and probability distributions (right panel) are shown. In the IF2 without lipid replica 1 (IF2_nolipid_rep1) simulation, a lipid molecule interacts at the TM4/TM6 interface after 400 ns. (**E**) Stability of the substrates in all simulated conditions.

### Bound lipids loosen drug-transporter interactions in the IF conformation

To explore the functional role of these lipids in the transport mechanism, we carried out MD simulations of BmrCD_IF2 in the presence and absence of bound lipids monitoring the dynamics of bound lipids, the stability of the substrates under all conditions, and the distance between TM4 and TM6 where the H184-R304 latch is located. Most of the 27 lipid molecules identified from the cryo-EM density were flexible, showing atomic fluctuations larger than 3.5 Å for the phosphorous atom of the headgroups, and even larger tail deviations. Only 6 molecules showed atomic fluctuations between 2.7 and 3.5 Å (Fig. S13) and these do not include the two lipid molecules near the latch. Indeed, lipids 4 and 23 appeared quite flexible.

We monitored the Cα-Cα distance for H184-R304 under all simulated conditions as illustrated by the time series and its probability distribution (Fig. 4D and Table S3). The analysis of the RH latch revealed that the presence of a single substrate molecule (Fig. 4D, gray) stabilizes shorter distances, while introducing a second substrate (silver) triggers the opening of the TM4/TM6 interface (shoulder in the silver distribution). The presence of two lipid molecules keeps the interface more open (extremely pronounced in replica 1 of the lipid bound simulation set, red curve), although we observed that it began to close as the lipids started to depart from the site during the simulation (red curve). Simulations after removal of the lipid bound molecules (green) show two peaks in the distance distributions. Remarkably, the interface begins to close until a new lipid molecule, from the bulk lipids, comes in after 400 ns (snapshots from the simulations are shown in Fig. S14A). The implied role of the TM4/TM6 interface as a gate is in agreement with previous structural studies of mammalian Pgp and MD simulations of the ABC heterodimer TM287/TM288(*23, 24*).

To monitor the substrate stability, we calculated the atomic fluctuations of all HT heavy atoms as time series for all simulated conditions (Figs. 4E and S14). Furthermore, to map the movement of the substrates during the simulations, we represented the HT molecules as vectors connecting the nitrogen of the piperazine to the ethoxy group (reported every 100 frames, Fig. 4E and S14). When only one HT molecule is present, it is free to explore the wide BmrCD cavity (Fig. 4E, gray curve). In contrast, when two HT molecules are bound (Fig. 4E, silver), the first one becomes more flexible and starts packing against the second one (as shown by the blue and magenta vectors which tend to cluster together in Fig. S14C). The presence of lipid molecules prevents substrate packing and pushes the first substrate towards the extracellular half of the protein at the apex of the vestibule (Fig. S14C). Once it pushes the substrate, the lipid molecule leaves the interface presumably allowing the transporter to switch from inward-open to inward-occluded prior to transitioning to the outward open conformation to extrude the substrates.

## DISCUSSION

The structures presented here paint an almost complete model of the conformational landscape of BmrCD at near atomic resolution (Fig. 5). In this model, the substrate-coupled cycle includes previously (*8*) described stepwise binding of Hoechst and Mg^2+^ and culminating with the population of the OF conformation reported here. B-factor analysis (Fig. S15) shows that the NBDs are dynamic in the IF state prior to formation of the catalytic NBSs, whereas the ECD becomes dynamic in the OC and OF states presumably to facilitate Hoechst release. Following dissociation of Hoechst, the transporter transitions to the OC conformation. Previous DEER analysis demonstrates that the population of the OF conformation in lipid nanodiscs in the ADP-Vi trapped state requires the binding of Hoechst (*8, 17*). The stability of the drug-bound OF conformation in BmrCD as opposed to other ABC exporters can be attributed among other factors to the ECD which may increase the energetic cost of drug release, and the smaller angle of extracellular opening relative to other ABC exporters that may promote stronger substrate interactions with the transporter in the OF state (Fig. S16 and Table. S4).

**Fig. 5.**
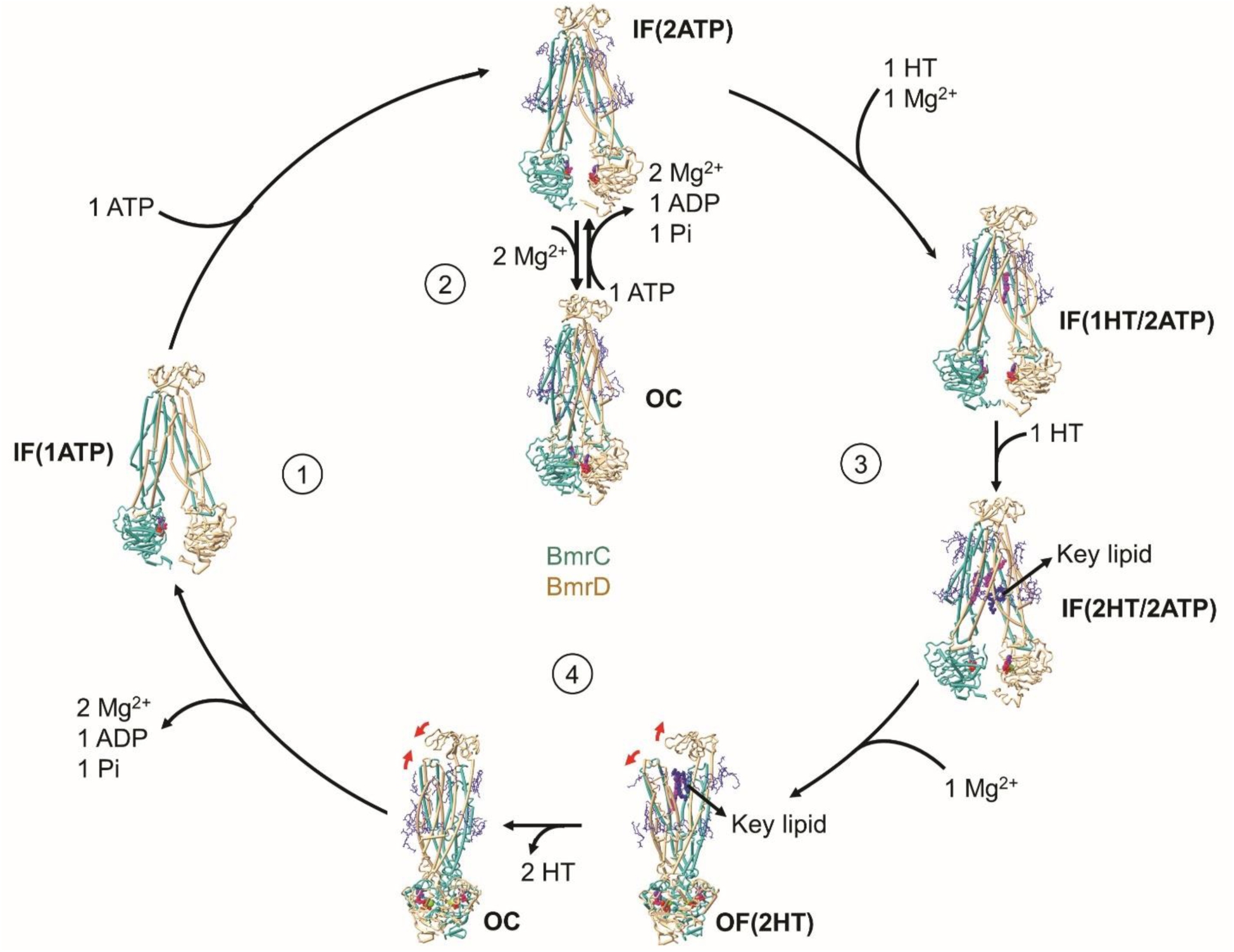
Working model for lipid-mediated substrate transport. BmrCD is shown in cartoon with helices represented by cylinders. Hoechsts, key lipids: lipids POV23 in IF(2HT/2ATP) and POV15 in OF(2HT), and ATP/ADPVi are shown by ball-and-stick model. The rest of the lipids are shown in sticks. Mg^2+^ ions are shown in green ball. Red arrows indicate the conformational changes. Four steps of the transport cycle are indicated by circled numbers. The structure models are as follows: IF(1ATP), AlphaFold3 generated model (See details in Materials and Methods); IF(2ATP), PDB 8FPF; IF(1HT/2ATP), PDB 8SZC; IF(2HT/2ATP), BmrCD_IF2; OF(2HT), BmrCD_OF; OC, PDB 8T1P.

Indeed, unique to our model is the profiling of interactions that stabilizes bound substrates to both IF and OF intermediates. Contacts between Hoechst and the two protomers are asymmetric with BmrC contributing the majority of interacting residues. This finding echoes the asymmetry that shapes the conformational cycle previously noted (*8, 21*) and reinforced here by the structure of the OF conformation. The BmrC side of the V-shaped extracellular vestibule is closer to the substrate (3.4 Å), display a more flexible TM1, and an opening near the exit. In contrast, the BmrD side is further from the substrate (8.9 Å), has a less flexible TM1, and an electrostatic interaction between K40^BmrD^ and R381^BmrD^ may obstruct the exit (Fig. S17).

Starting from lipid molecules bound at the entrance of the OF vestibule, the MD simulations reveal how lipids act to disrupt the interactions of the Hoechst molecules with the transporter. While the driving force for movement of the substrate from the IF to the OF vestibule is contributed overwhelmingly by the energy of ATP binding and hydrolysis that power conformational changes, a mechanism is needed for the release from the OF conformation. Functional and spectroscopic data of BmrCD harmonize with our model (*8, 17, 21*). Formation of the OF conformation for BmrCD in lipids requires Hoechst binding. In addition, the OF stability is reduced upon transition from detergent micelles to lipid bilayers(*8, 17, 21*).

The energetic cost of substrate release is dependent on the strength of its interactions with the OF conformation. It is possible that despite the higher concentration of substrate on the release side of the membrane, its intrinsic affinity is weak enough such that its dissociation is spontaneous. Such spontaneous dissociation was previously observed in MD simulations of TM287/TM288 (*23*) and was ascribed to changes in hydration. Alternatively, for those transporters wherein ATP hydrolysis occurs after the formation of the OF conformation, the energetic drive to reset to the IF conformation could also overcome low-affinity binding of the substrate to the OF state.

These models, however, do not explicitly include the energetic contribution of lipids. The amphipathic substrates of multidrug ABC transporters tend to be soluble in the lipid phase. Indeed, early models of ABC transporters posited a hydrophobic “vacuum cleaner” model where substrate molecules are bound from the inner leaflet of the membrane and extruded into the outer leaflet (*25*). Conversely, lipids are amphipathic, and their high concentrations can overcome their intrinsic low affinity to the transporters and their entropy, thereby driving binding to the transporter. The MD simulations establish that lipids loosen stabilizing interactions between the transporter and substrate prior to release.

Thus, we propose a dynamic “competition” model between substrates and lipids for multidrug ABC exporters. We posit that this competition results from the modest affinity of the substrates to these polyspecific transporters and the high concentration of lipids which have access to the rather wide vestibules in both IF and OF conformations. The constant entry and exit of lipid molecules disrupt interactions with bound substrates. It is supported by previous MD simulations for other ABC transporters in the IF conformations where lipids spontaneously enter the vestibule (*24, 26*). A lipid-induced modulation of substrate affinity may be a general mechanism for other ABC exporters that are more specific to amphipathic substrates.

## MATERIALS AND METHODS

### BmrCD cloning, expression, and purification

Wild-type BmrCD was cloned in the pET21b(+) plasmid and then transformed into *Escherichia* coli BL21(DE3) cells to be expressed in minimal media supplemented with 0.5% v/v glycerol, 2.5 μg/ml thiamin, 100 μg/ml ampicillin, 1mM MgSO4, and 50× MEM amino acids. Cell cultures were grown at 37 °C until the optical density at 600 nm (OD600) reached 1.2. Subsequently, induction of BmrCD expression at 25 °C was initiated by adding 0.7 mM isopropyl β-d-1-thiogalactopyranoside (IPTG). Cells were harvested after 5 hours of expression and stored at -80 °C till purification.

For BmrCD purification, cell pellets were resuspended in a lysis buffer containing 50 mM Tris-HCl (pH 8.0), 1 mM EDTA, 1 mM PMSF, and a complete EDTA-free protease inhibitor cocktail tablet (Roche) before undergoing sonication. The lysate was centrifuged at 9000 x g for 10 minutes at 4 °C, yielding the supernatant utilized for high-speed centrifugation at 185,000 x g for 1 hour at 4 °C to precipitate membrane pellets. To solubilize the membrane pellets, a resuspension buffer comprising 50 mM Tris-HCl (pH 8.0), 0.1 M NaCl, 15% v/v glycerol, 1 mM DTT, and 1.25% w/v n-dodecyl-β-D-maltopyranoside (β-DDM) was applied for 1 hour on ice. Following this, centrifugation at 185,000 x g for 1 hour at 4 °C separated insoluble material, and the supernatant was combined with pre-equilibrated Ni-NTA beads for a 2-hour incubation in binding buffer (50 mM Tris-HCl, pH 8.0, 0.1 M NaCl, 15% v/v glycerol, 0.05% β-DDM). After resin washing with binding buffer supplemented with 20 mM imidazole, protein elution was performed using binding buffer containing 250 mM imidazole.

Subsequently, protein purification involved size exclusion chromatography (SEC) with a Superdex 200 Increase 10/300 column (Cytiva). The SEC buffer 1 which consisted of 50 mM Tris-HCl (pH 7.5), 0.15 M NaCl, 10% v/v glycerol, and 0.05% β-DDM was used for equilibration.

### BmrCD nanodiscs preparation

The preparation of BmrCD nanodiscs followed the procedure outlined previously(*8*). In summary, lipids consisting of PC (L-α phosphatidylcholine) (Cat #: 840051, Avanti Polar Lipids, Alabaster, USA) and PA (L-α phosphatidic acid) (Cat #: 840101, Avanti Polar Lipids, Alabaster, USA) were combined in a molar ratio of 9:1. The BmrCD nanodiscs were then created by mixing lipids, MSP1D1, and BmrCD micelles at a molar ratio of 1800:360:3:1 (β-DDM: lipid: MSP: BmrCD). The mixtures were incubated for 30 minutes before adding biobeads (0.8-1 g/ml) and rotating at 4 °C overnight. Following the removal of biobeads by centrifugation, BmrCD in nanodiscs was filtered through a 0.45 μm filter and subsequently purified via size exclusion chromatography using a Superdex 200 Increase 10/300 column (Cytiva) with SEC buffer 2 (50 mM Tris-HCl, pH 8.0, 0.15 M NaCl).

### Cryo-EM sample preparation and data acquisition

Reconstituted BmrCD at a concentration of approximately 3 mg/ml was incubated at 37 °C for 15 minutes with a mixture containing 10 mM ATP, 4 mM MgSO4, 4 mM vanadate, and 1.2 mM Hoechst-33342 (Thermo Fisher Scientific) before plunging. Cryo-EM grids were prepared by applying 2.5 μl of the sample to a Quantifoil UltrAuFoil grid (R1.2/1.3, 300 mesh, Electron Microscopy Sciences) using a Vitrobot Mark IV (Thermo Fisher), pre-equilibrated at 4 °C with 100% humidity. Grids suitable for cryo-EM data collection were selected through screening by the Glacios system (Thermo Fisher Scientific).

Cryo-EM data collection was conducted using a 300 keV Titan Krios G4 microscope (Thermo Fisher Scientific) equipped with a Gatan K3 direct electron camera. Movies containing 50 frames were recorded with three exposures per hole using EPU in normal mode. The magnification was set at 130,000 with a physical pixel size of 0.647 Å/pixel, and defocus values were within the range of -0.9 to -2.0 μm. The total electron dose over the sample during data collection was 56 e-/Å².

### Cryo-EM data processing

The cryo-EM dataset was processed in cryoSPARC(*27*), implemented with MotionCor2(*28*) for beam-induced motion correction and Gtcf(*29*) for CTF estimations. For BmrCD_OF, BmrCD_OC was used as template for Template picker to identify approximately 6.6 million particles, followed by 2D classification, heterogeneous refinement, and Ab initio to obtain high-quality particles. Followed by obtaining the particle images for occluded conformation, one round of 3D classification was performed to get 27,802 out of 185,953 particles to generate the outward facing conformation map (Fig. S2). Subsequently, non-uniform(*30*) and local refinement were applied, resulting in a final map with a resolution of 3.11 Å. For BmrCD_IF1 and IF2, similar strategies were applied (See details in Figs. S10, S11 and S12).

All structural models refinement and validation were performed using Real-space refinement(*31*) and MolProbity(*32*) implemented in Phenix 1.21.1-5286(*33*). Coot(*34*) was applied to build the models. The predicted structure model of BmrCD_IF(1ATP) was generated by AlphaFold 3(*35*) with input files of the sequences of BmrC and BmrD, as well as 1 copy of ATP in autoseed mode. Visualization of the results and preparation of figures were carried out using UCSF Chimera 1.18(*36*), ChimeraX 1.4(*37–39*), and PyMol 2.4.1. Additionally, the B-factor was calculated using the B-factor putty tool implemented in PyMol 2.4.1 with default settings. solvent-accessible volume calculations were conducted using CASTp(*40*). Radius measurement of the vestibule was performed using the software HOLE(*41*).

### Molecular dynamics simulations

#### 1 Molecular dynamics simulation for BmrCD_OF

##### 1.1 BmrCD_OF model preparation

Using the cryo-EM input model BmrCD_OF, we first completed partially resolved lipids. To do this we used the PSFGEN utility in VMD(*42*) using the CHARMM36 forcefield(*43, 44*). Protein-lipid clashes were resolved by a brief energy minimization of 2,000 steps using NAMD3(*45*). The D141A mutation was modeled using mutate.tcl in VMD(*42*). The resulting wild-type (WT) and mutant models were then embedded in membranes generated with CHARMM-GUI(*46*). Each membrane consisted of a 9:1 ratio of PC:PA in accordance with the nanodisc composition used in our experiments and was solvated with TIP3P(*47*) water and ionized in VMD(*42*).

##### 1.2 Equilibrium molecular dynamics

All molecular dynamics (MD) simulations of the BmrCD OF state were performed using the NAMD3 engine(*45*). A standard equilibration protocol included the following scheme: (i) 10,000 steps of energy minimization. (ii) Gentle heating ramping up the temperature by 25K at a time for 1,000 steps of MD until the final temperature of 310 K was reached. (iii) 2.5 ns of equilibration with harmonic restraints (k = 1.5 kcal mol^-1^ Å^-2^) on lipid headgroups, protein backbone, Hoechst molecules, ATP and coordinating Mg^2+^ ions. (iv) 2.5 ns of equilibration with harmonic restraints (κ = 1.0 kcal/mol/Å^2^) on protein backbone, Hoechst molecules, ATP and coordinating Mg^2+^ ions. (v) 2.5 ns of equilibration with harmonic restraints (κ = 0.5 kcal/mol/Å^2^) on protein backbone and Hoechst molecules. (vi) 5.0 ns of unrestrained equilibration.

After equilibration, four replicates for WT and three replicates for mutant were simulated for 2 µs resulting in an aggregate of 14 µs simulation data. Simulation parameters used included a 12- Å cutoff distance for long-range interactions, a 10-Å switching distance, a 2-fs timestep, the particle mesh Ewald (PME) method for electrostatics(*48*), Nosé-Hoover Langevin piston for pressure coupling with a target of 1 atm, oscillation period of 100 fs and decay of 50 fs, and Langevin dynamics for temperature control with a damping coefficient of 1 ps^-1^(*49*).

##### 1.3 Analysis

To investigate the influence of lipids on HT binding and release, we first applied kernel density estimation (KDE) to map the density distribution of lipid molecules, thereby identifying high-occupancy regions accessible to phospholipids. Building on this initial analysis, we quantified native contacts and non-bonded interaction energies between HT and BmrCD using the NAMD Interaction Energy module in VMD(*42, 45*), which allowed us to assess lipid-induced perturbations in substrate binding. Furthermore, we analyzed distribution of HT tilt angle and changes in salt bridges to elucidate lipid-induced conformational shifts. As our investigation deepened, we evaluated the structural dynamics by measuring the backbone root-mean-square deviation (RMSD) of BmrCD transmembrane helices over time, revealing rearrangements associated with lipid binding and substrate displacement. Additionally, we quantified lipid– substrate interactions by tracking the number of lipid heavy atoms within 3.5 Å of HT. Finally, we calculated the minimum distance between E51 and E75 of BmrD and HT over time, as well as the distribution of extracellular domain (ECD) domain angles relative to the negative z axis (0, 0, -1).

To determine the binding stability of HT-1 and HT-2 molecules in the binding pocket of BmrCD, we calculated the RMSD of each molecule and averaged it over the three replicates of D141A and WT, to compare them quantitatively. The first frame of the trajectory for either structure was selected as the reference. Each frame of the trajectory was aligned to the reference structure to remove the translational and rotational motions. To quantify ionic interactions, ion pairs between positive or negative residues and the ligands within 5.0 Å, were considered. All analyses were performed using Python 3.9 Plots were generated with Matplotlib to visualize interaction patterns. MDAnalysis 2.8.0 was used to identify molecular contacts based on distance criteria(*50*). Analysis was performed using custom scripts in VMD(*42*) or Python using the following packages: SciPy(*51*), MDAnalysis(*50*), and Seaborn(*52*).

#### 2 Molecular dynamics simulations for BmrCD_IF1 and BmrCD_IF2

##### 2.1 BmrCD IF models preparation

Three distinct BmrCD protein structures were employed here, specifically pdb IDs 8FMV (BmrCD_IF) and 8SZC (BmrCD_1HT/ATP)(*8*), alongside a newly solved structure, BmrCD_IF2. These structures, chosen for their relevance to the BmrCD transporter mechanism, included variations in ligand binding—two structures with dual Hoechst (HT) molecules and one with a single HT molecule. BmrCD_IF2 revealed bound lipid molecules, especially at the interface between transmembrane helices 4 and 6 (TM4/6). To elucidate the role of this lipid, two different simulation setups were prepared: one including the 27 bound lipids (BmrCD_IF2) and one without them (BmrCD_IF2_nolipid) (Table S2).

The Molecular Operating Environment (MOE) software, v2022.02 (*53*) was utilized to model missing loops for the protein and missing heavy atoms for the solved lipid molecule. Hydrogens were added according to the physiological pH (7.4).

##### 2.2 System Preparation and Molecular Dynamics Simulation

The Amber22 software (*54*) was used to prepare each system. The *packmol-memgen* tool facilitated membrane embedding within a lipid bilayer composed of 9:1 PC/PA, mirroring the experimental lipid composition. The systems were solvated in a 150 mM NaCl solution, with topologies and initial coordinates generated via *tleap* (*54*) using the *amber19SB* force field for the protein, *lipid21* for the lipid and the OPC water model. All structures were simulated with ATP and Mg^2+^ bound at the consensus site, and ATP alone at the degenerate site. The ATP topologies were sourced from the Bryce group’s website, utilizing parameter sets detailed by Meagher et al (*55*).

##### 2.3 Ligand Preparation

The Hoechst molecule was computationally refined using Gaussian 16(*56*) to ensure accurate molecular interactions in subsequent simulations. Geometry optimization was first performed with the B3LYP functional combined with the 6-31G* basis set(*57, 58*), enforcing a stringent (*tight*) convergence criterion on the self-consistent field (SCF) calculations. Following this step, the electrostatic potential (ESP) was calculated using the Hartree-Fock method with a 6-31G* basis set, coupled with Merz-Kollman population analysis to accurately determine the electrostatic potential surface. Subsequently, the Amber22 *Antechamber* tool (*54*) was employed to fit RESP charges to the optimized structure, which were then utilized in molecular dynamics simulations to accurately model HT’s behavior.

##### 2.4 Molecular Dynamics Protocols

Each prepared system was subjected to an initial energy minimization to relieve any steric clashes or unfavorable geometries, followed by a canonical ensemble (NVT) heating step where the system was gradually heated from 0 to 310 K over 200 ps, and then sustained at this target temperature for an additional 50 ps. The NVT phase employed a 1 fs timestep with positional restraints applied to all heavy atoms, excluding water oxygens and ions in solution. Upon reaching the desired temperature, the systems were subjected to the isothermal-isobaric (NPT) equilibration phase to stabilize pressure and density, using semi-isotropic pressure scaling. The positional restraints on proteins, ligands (inclusive of ATP, Mg^2+^, and HT molecules), and lipid head groups were gradually released over a period of 10 ns. This stepwise easing of restraints—from 10.0 kcal/mol/Å^2^ applied to the heavy atoms of the protein and ligands and 2.5 kcal/mol/Å^2^ applied to the lipid head groups—culminated in the total removal of restraints for the last equilibration step prior to the NPT production run of ∼ 4 ms. Such a gradual approach facilitated orderly water diffusion, membrane equilibration, and stabilization of protein binding sites and ligands. Temperature regulation was managed using the Langevin dynamics algorithm(*59*) stabilizing the system at 310K. The pressure was controlled at 1 bar using the Monte Carlo Barostat (*60*), which operates with a friction coefficient of 1 ps^-1^. The timestep for the simulations was set at 1 fs during the initial NVT and first NPT equilibration steps and was increased to 2 fs for the remainder of the equilibration stages and the production phase. The SHAKE algorithm (*61*) was used to constrain hydrogen atoms, ensuring the stability of each system, and allowing for a larger timestep. Long-range electrostatics interactions were calculated via the particle mesh Ewald method, while the cutoff for van der Waals and Coulombic interactions was set between 10 and 12 Å to ensure accurate force field representation.

Analysis were performed using *cpptraj* from the *AmberTools23* package suite or custom scripts in VMD (*42*) using the tcl scripting language. Plots were generated using custom python scripts and matplotlib.

## Data availability

The cryo-EM maps of BmrCD_OF, BmrCD_IF1, and BmrCD_IF2 have been deposited in the Electron Microscopy Data Bank (EMDB) under accession codes EMD-45940, EMD-45938, EMD-45939, respectively. The atomic coordinates of BmrCD_OF, BmrCD_IF1, and BmrCD_IF2 have been deposited in the Protein Data Bank (PDB) under accession codes 9CUS, 9CUP, and 9CUR, respectively. For MD simulation, initial coordinates, simulation input files and coordinate files of the final output are provided as Data S1 and S2.

## Acknowledgments

We acknowledge the use of the Glacios cryo-TEM for grids screening, and the cryo-EM data were collected at the Center for the Structural Biology Cryo-EM Facility at Vanderbilt University. We appreciate Erkan Karakas for his help in structure analysis.

## Author contributions

Q.T. performed all the experiments. M.S. and Y.Z. performed the MD simulations for BmrCD_OF under the supervision of E.T. P.B. performed the MD simulations for BmrCD_IF2. Q.T. and H.S.M. designed the research, analyzed, and interpreted the structures and wrote the manuscript with input from all the authors.

## Supplementary Materials

**Fig. S1.**
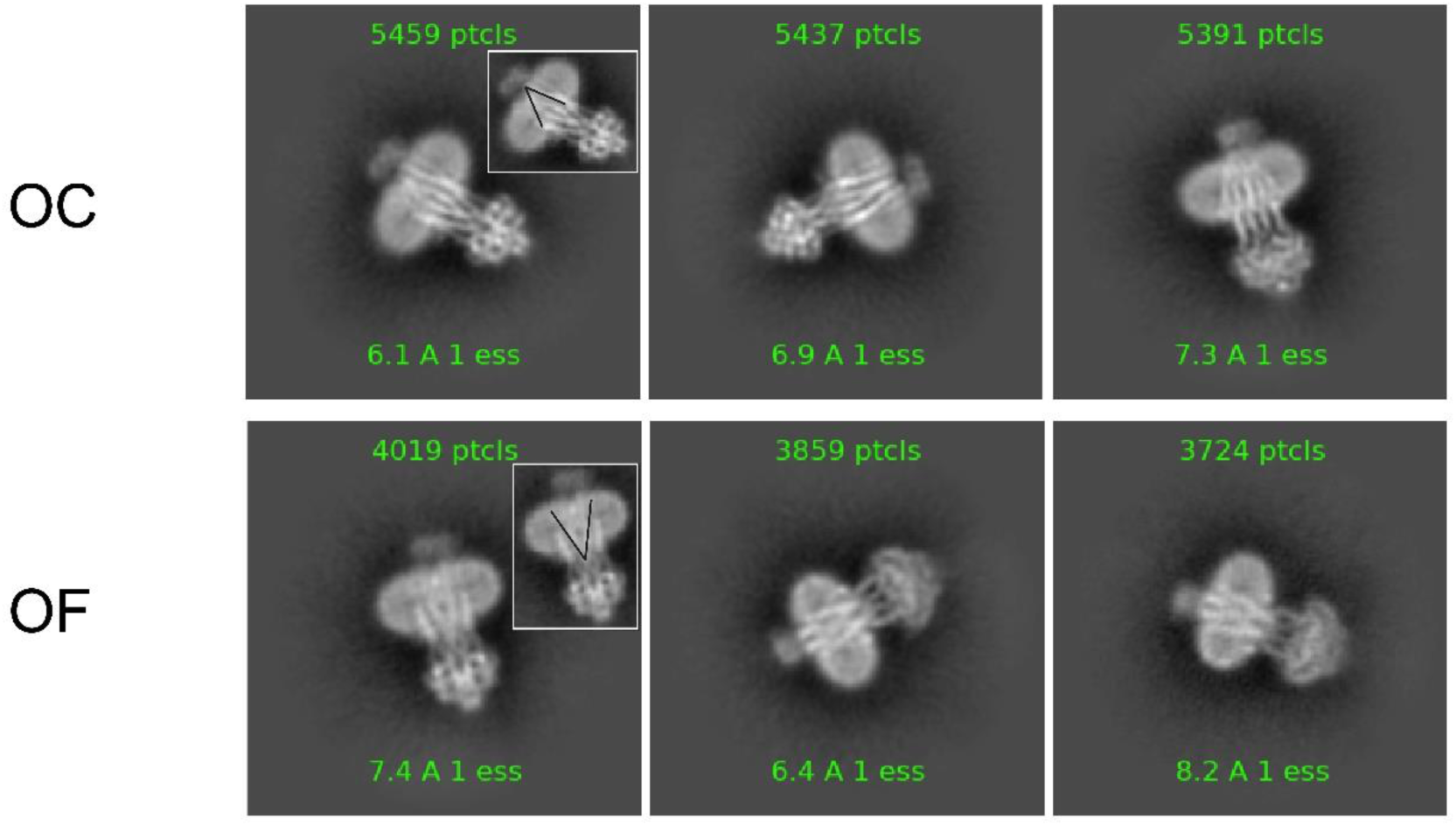
Representative 2D images for OC and OF conformations. The opening of the intracellular side for occluded (OC) conformation and the opening of extracellular side for outward-facing (OF) conformation are depicted as angle in the insets, respectively. The box size is 480 Å.

**Fig. S2.**
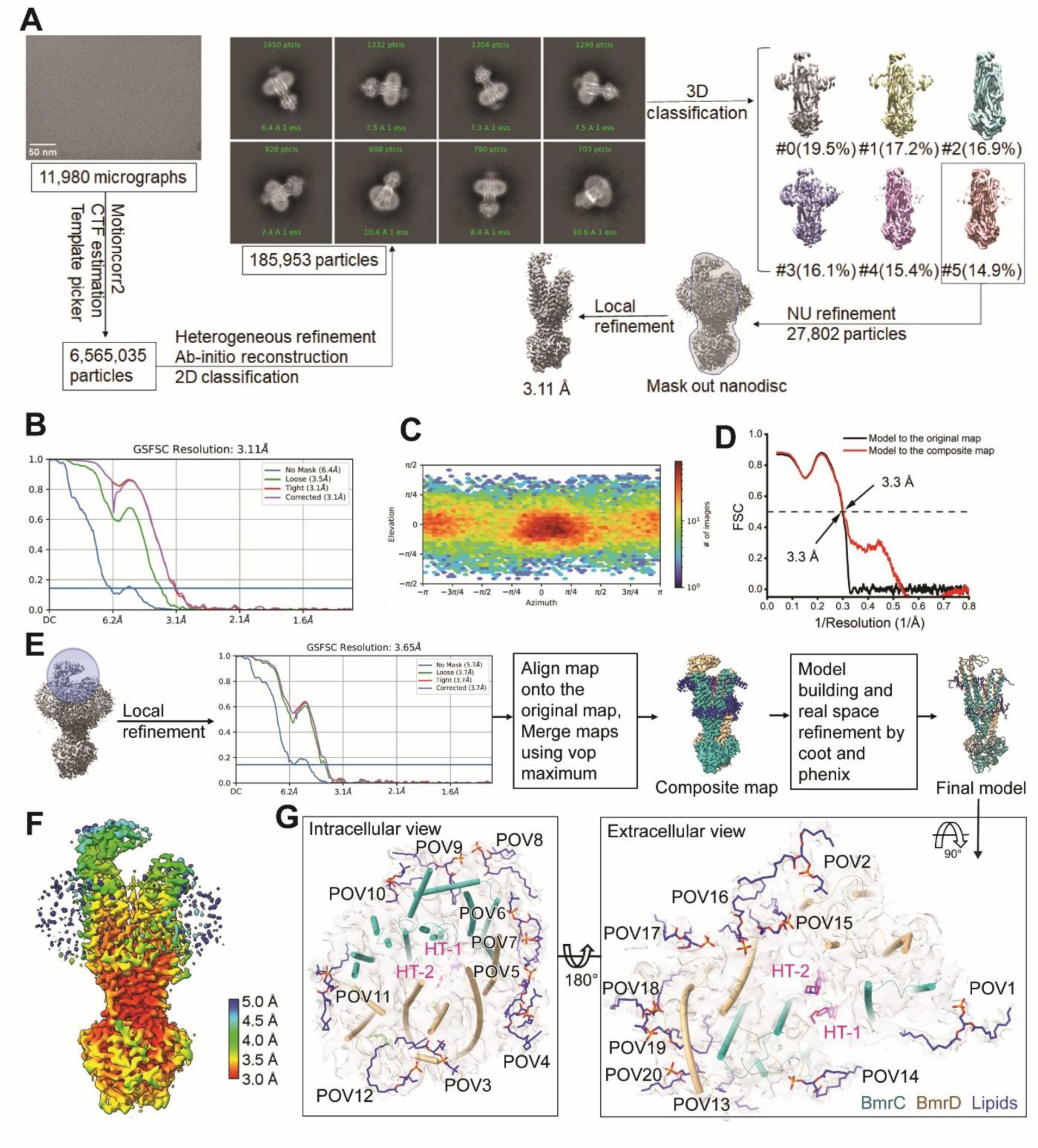
Cryo-EM data processing for BmrCD_OF. (**A**) CryoSPARC work flow for BmrCD_OF with representative micrograph. 2D and 3D images are shown. (**B**) Fourier shell correlation (FSC) curves for original map after local refinement in cryoSPARC. (**C**) Angular distribution of particles for original map after local refinement in cryoSPARC. (**D**) Fourier shell correlation (FSC) curves of the refined model versus the original map after Non-uniform refinement in cryoSPARC (black) and the composite map (red) are plotted. (**E**) Local refinemnet strategy for BmrCD_OF. Mask is shown in transparent cycle. (**F**) Local resolution map for original map after local refinement in cryoSPARC. (**G**) Lipids bound to the V-shaped interface of BmrC and BmrD. The map is shown as a transparent surface, with Hoechsts and lipids highlighted.

**Fig. S3.**
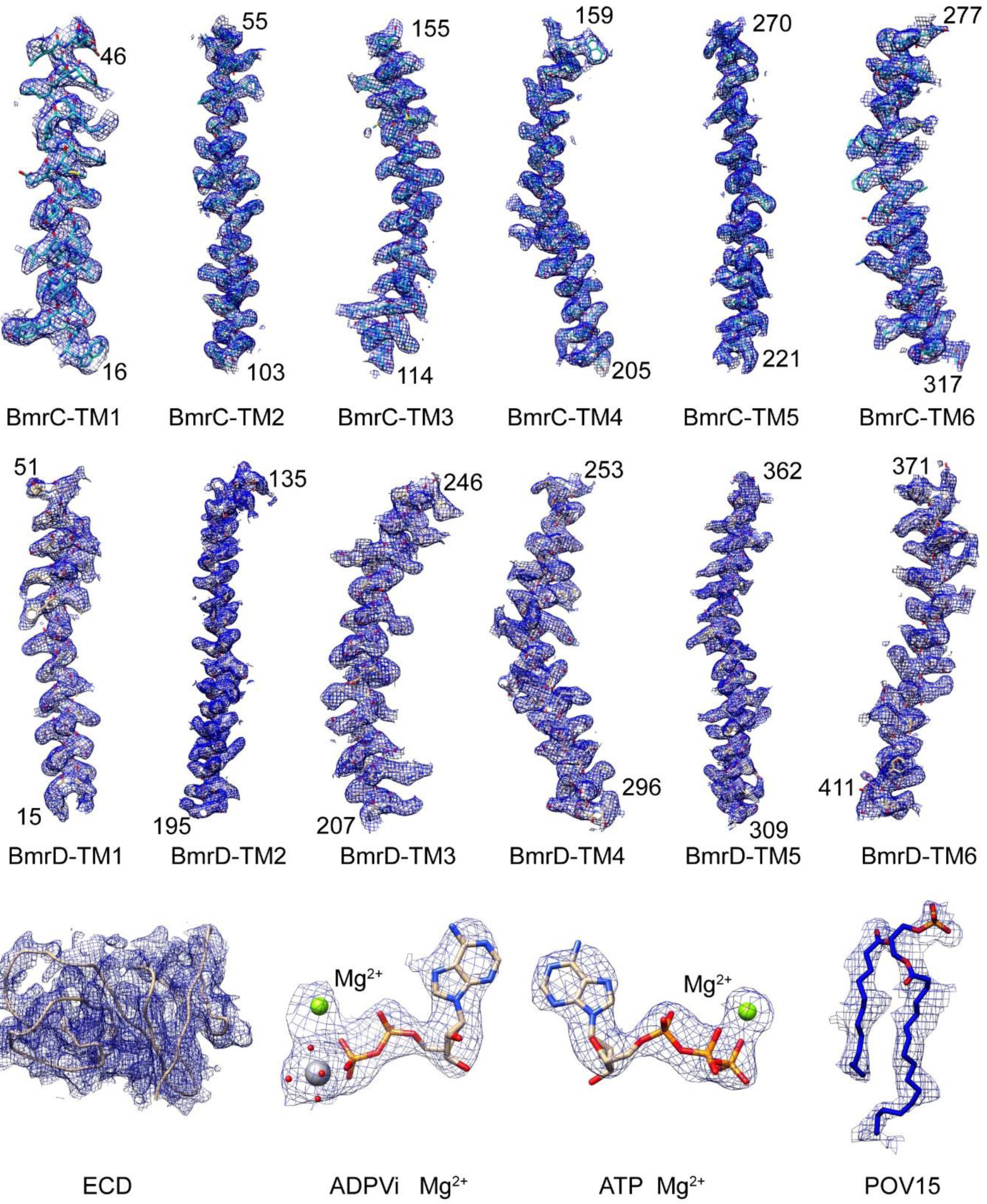
Cryo-EM densities for BmrCD_OF. Representative cryo-EM densities: transmembrane helices of BmrC and BmrD, extracellular domain (ECD), ADPVi, ATP, and a lipid molecule (POV15). The residue ranges for the helices are labeled.

**Fig. S4.**
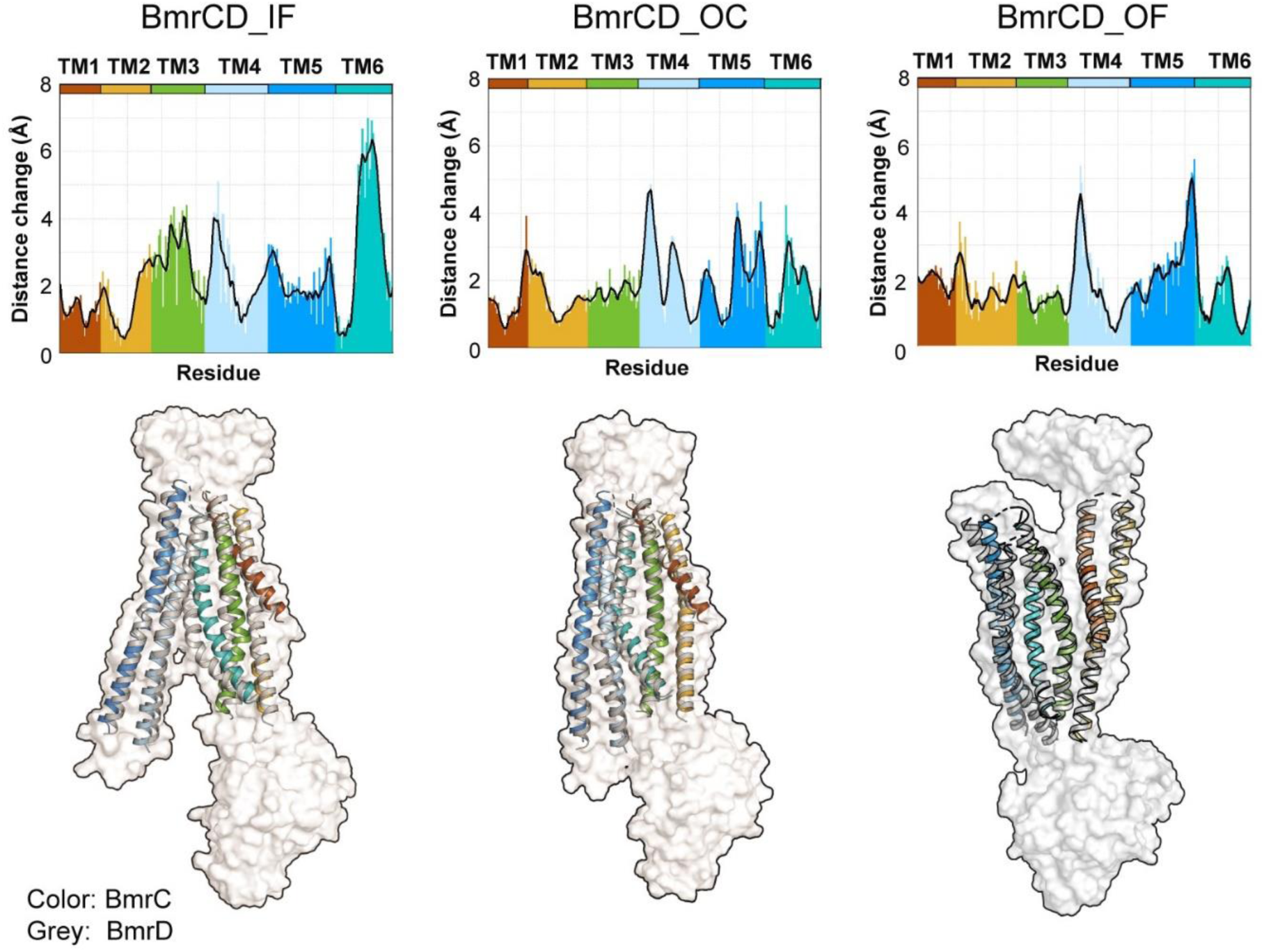
The asymmetry of TMDs among different BmrCD conformations. Upper panels: Distances between alpha cardons in the TMDs of BmrC and BmrD calculated by TMalign(*62*) after superposition of the two TMDs. Lower panels; cartoon representation of the superimposed TM helices from BmrC (colored according to the upper panels) and BmrD (shown in grey). A transparent surface representation of BmrD is shown as reference. BmrCD_IF (PDB ID: 8FMV) and BmrCD_OC (PDB ID: 8FHK) are shown for comaprsion(*8*).

**Fig. S5.**
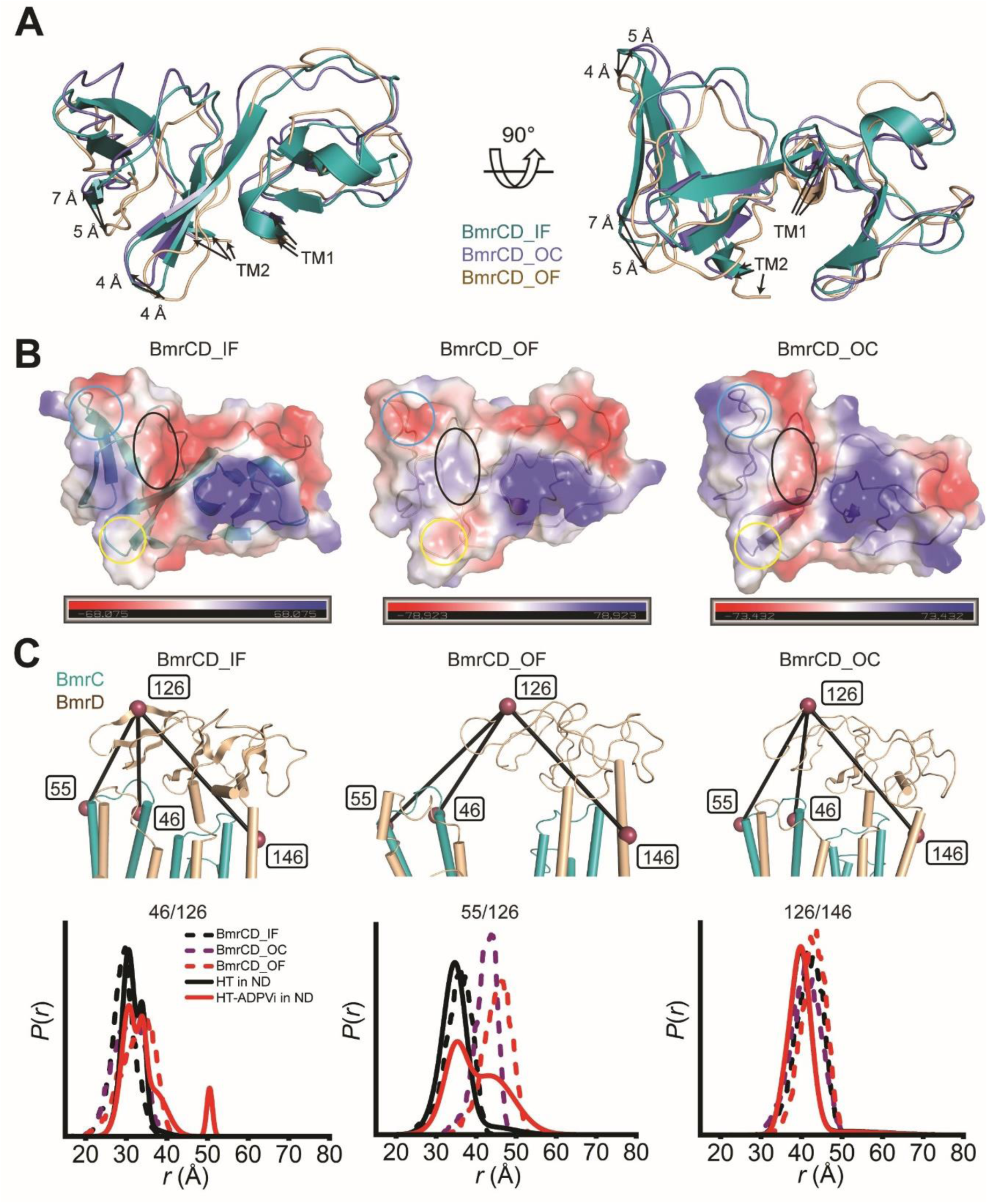
The ECD conformational changes. (**A**) Superimposition of extracellular domain (ECD) based on the alignment of BmrD in the three different conformations of BmrCD cryo-EM structures: BmrCD_IF (PDB 8FMV), BmrCD_OC (PDB 8T1P), and BmrCD_OF. The movement of the loops is highlighted. (**B**) Electrostatics potential surfaces of ECD in BmrCD cryo-EM structures are shown from bottom-up view. The color scale ranges from blue (positively charged regions) to red (negatively charged regions) whereas neutral regions are shown as white. The circles highlight the major changes among the IF, OC, and OF conformations. (**C**) DEER distance distributions for spin-label pairs highlighting the movement of the ECD. Upper panels: close-up views of the spin pairs between ECD and TMD in the IF, OF and OC conformations. Lower panels: experimental (solid lines) and simulated (dash lines) DEER distance distributions highlighting the movement of the ECD. The simulated distance distribution derived from cryo-EM structures by MDDS. 46/126, 55/126, and 126/146 refer to the spin label pairs for 46^BmrC^/126^BmrD^, 55^BmrC^/126^BmrD^, and 126^BmrD^/146^BmrD^, respectively. Color scheme is the same as Fig. 1C.

**Fig. S6.**
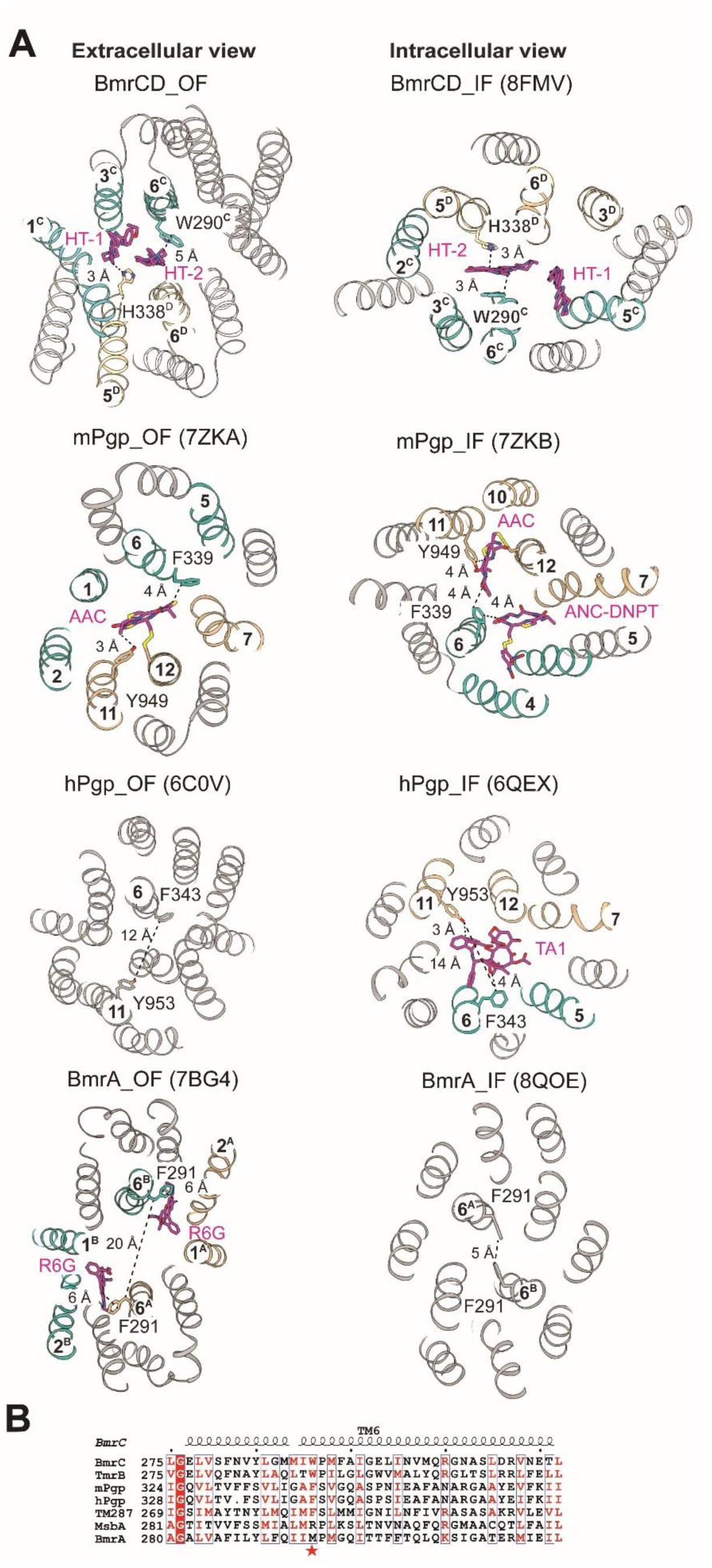
Equivalent WH-latch. (**A**) The extracellular views of the OF conformations (left panels) and Intracellular views of the IF conformations (right panels) highlight WH-latch in BmrCD and the equivalent latch in mouse P-glycoprotein (mPgp), human P-glycoprotein (hPgp), and F291 in BmrA homodimer. The protein is depicted in cartoon representation, with the side chains of the latch resides and substates shown as sticks. The substate-binding helices from BmrC, the N-terminal of mPgp and hPgp, subunit B of BmrA are shown in light sea green, while those from BmrD, C-terminal of mPgp and hPgp, subunit A of BmrA are depicted in tan; the remaining parts are in grey. Helices number and the distances are indicated. PDB IDs(*8, 10, 12, 15, 16, 22*) are listed in parentheses. F291-F291 latch is closed in BmrA_IF without substrate. The distance of F343-Y953 latch of hPgp_OF without substrate is shorter than that of hPgp-IF with substrate. (**B**) Sequence alignment reveals that the W290 is conserved among heterodimeric ABC transporters, including TmrAB, TM287/288, and Pgp, but not conserved in homodimeric transporters such as MsbA and BmrA.

**Fig. S7.**
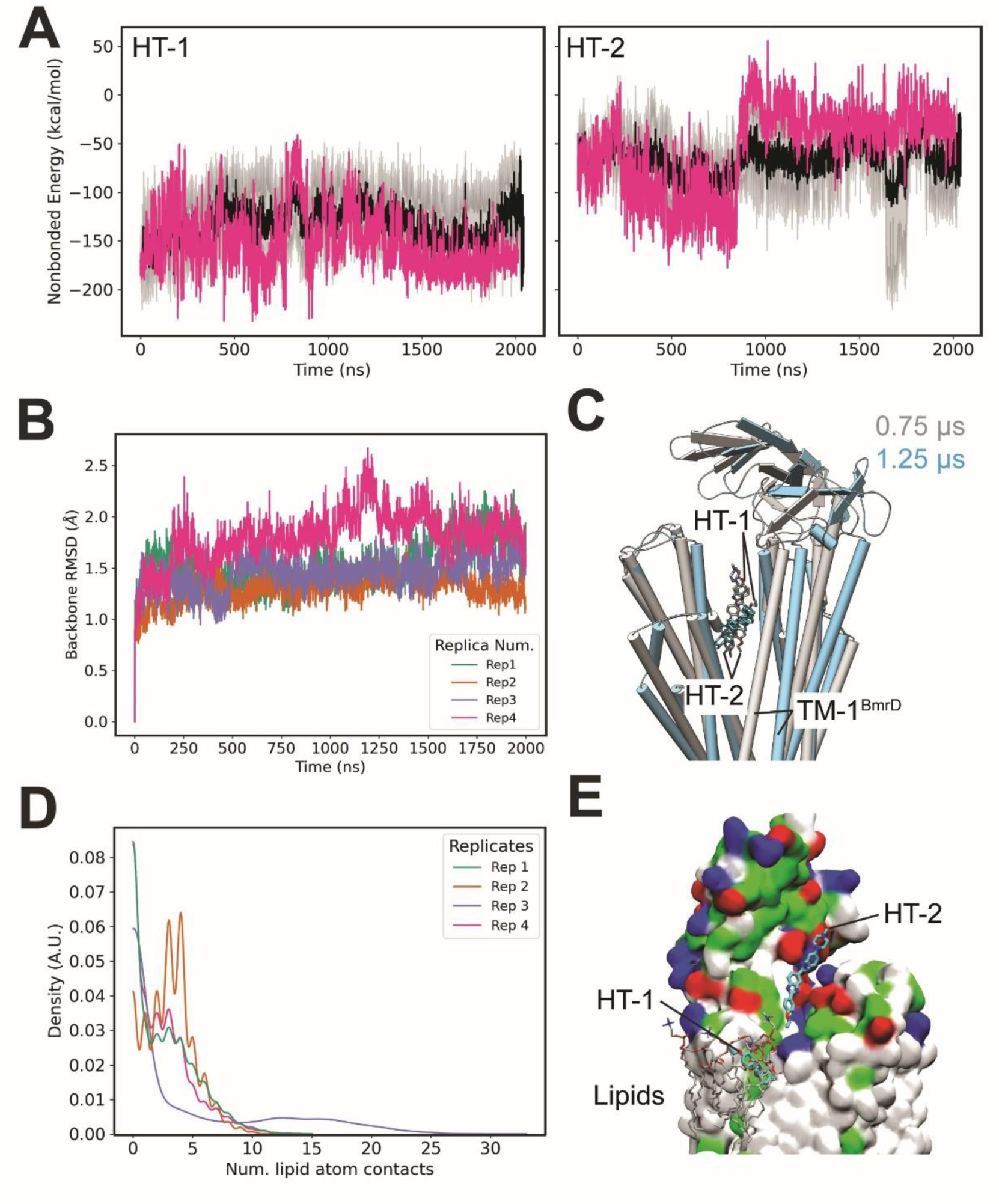
MD simulations of BmrCD_OF (part. **I)**. (**A**) Time series of non-bonded interaction energy between protein and each Hoechst molecule computed using the NAMD Interaction Energy module of VMD. (**B**) Time series of helical RMSD, measured using backbone atoms of transmembrane helices. Each trace represents an independent replica simulation. (**C**) Molecular rendering depicting the shift of Hoechst from 0.75 µs to 1.25 µs in grey and cyan, respectively. Note the shift of BmrD TM 1 as Hoechst is pushed out of its bound state. (**D**)Hoechsts-lipid contacts. Units are number of lipid heavy atoms within 3.5Å of Hoechst heavy atoms. (**E**) Molecular rendering of lipids (from the bulk membrane) surrounding HT-1, and the interaction of the ECD domain with HT-2. HTs and lipids are shown as cyan and grey sticks colored by heteroatom. Hydrophobic residues are marked as white, polar as green, basic as blue and acidic as red.

**Fig. S8.**
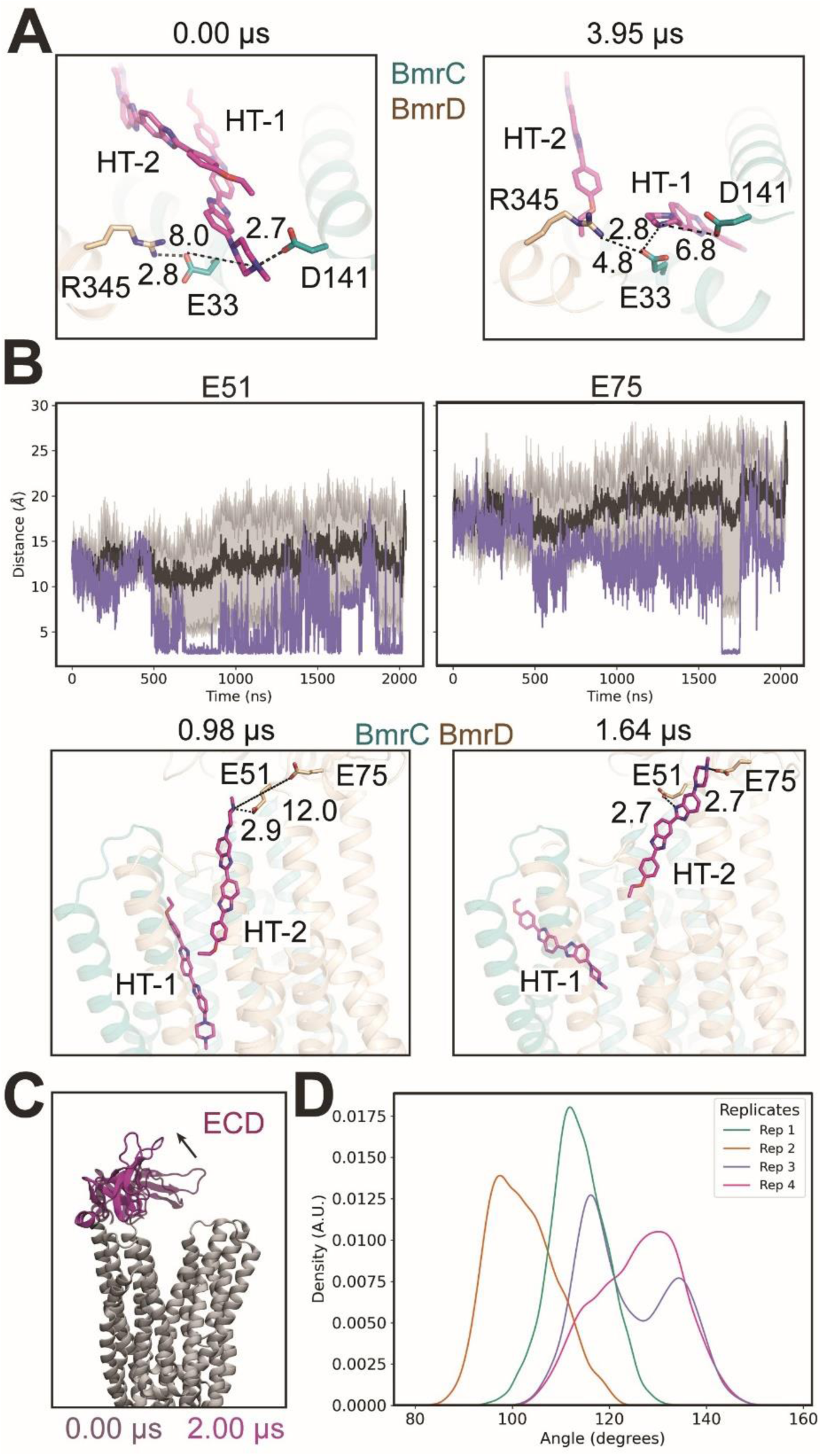
MD simulations of BmrCD_OF (part II). (**A**) Molecular rendering showing E33 (BmrC) interaction network changes. Distances are in Å. (**B**) Minimum distance (Å) measured between E51 and E75 of BmrD and Hoechst over time. The black trace depicts the mean distance over all trajectories with the grey area corresponding to the standard deviation. Purple trace shows the distance for replica 3 in which these interactions are observed. As E51-Hoechst interaction dissipates, E75-starts interacting with Hoechst. The molecular renderings shown below. (**C**) Molecular rendering showing the initial (transparent, 0.00 µs) and final (opaque, 2.00 µs) positions of the structural ECD domain from replica 3 in which Hoechst is observed to be interacting with it. The arrow indicates the movement. (**D**) Distribution of ECD domain angles as measured in reference to the negative Z-axis vector (0 0 1).

**Fig. S9.**
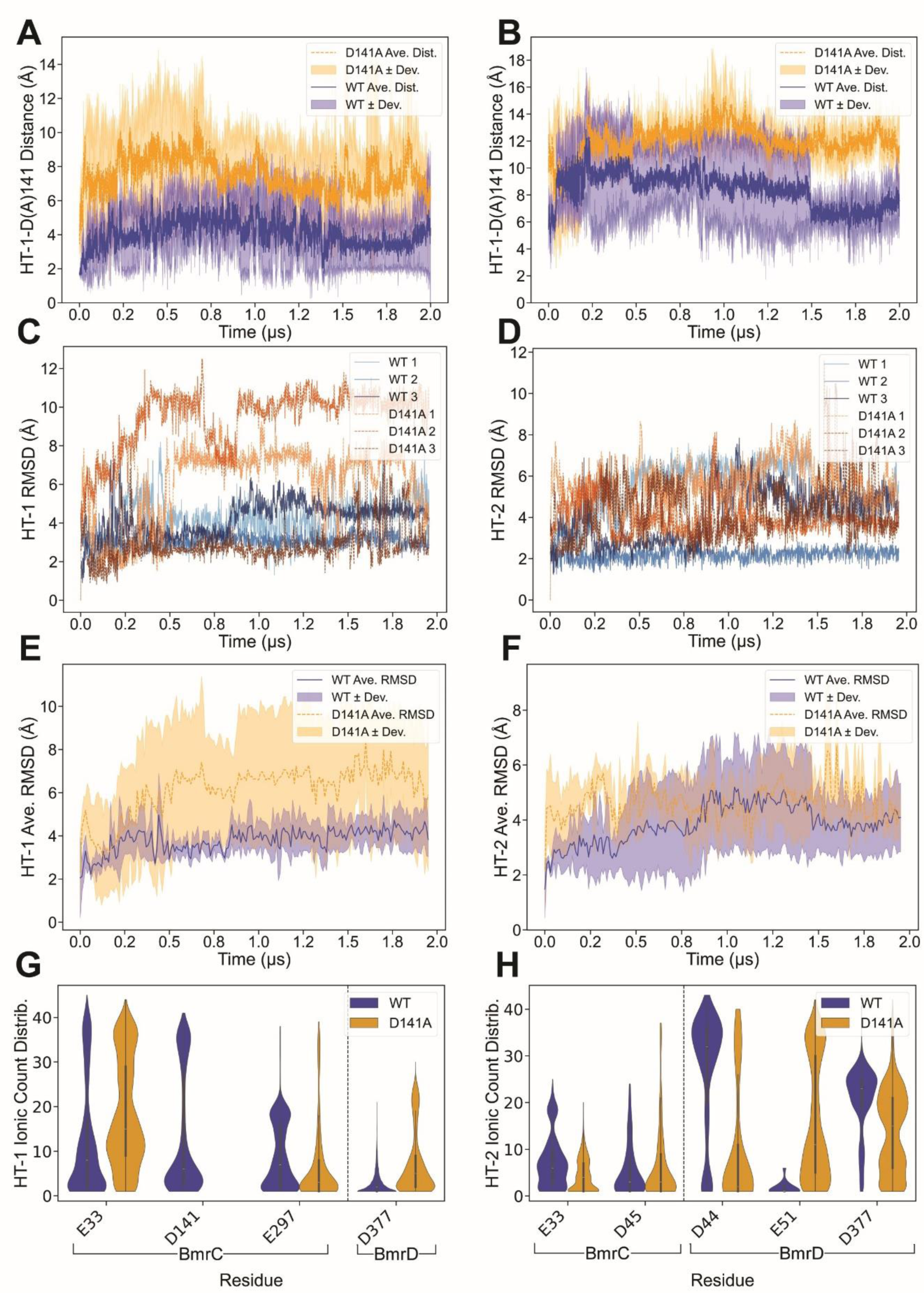
Distance evolution, RMSD and ionic bond counts for HT-1/HT-2 in wild-type (WT) vs. mutant (D141A). (**A**, **B**) The distance between COM of HT-1 or HT-2 and COM of residue D141 or D141A over the simulation. WT (blue) maintains shorter distances to D141, whereas D141A (orange) exhibits larger distances to the substituted site. The consistently larger distances in D141A are in support of weakening the ligand interactions with the protein, allowing HT-1 or HT-2 to drift further from the binding region. (**C**, **D**) The time evolution of HT-1 or HT-2 RMSD for WT (blue) vs. D141A (orange). The WT traces generally show lower or more stable RMSD values (3.8 ± 1.0 Å), while the D141A traces display higher RMSD, underscoring the destabilizing impact of losing D141 (6.1± 3.1 Å). (**E**, **F**) Average RMSD of HT-1 or HT-2 over WT and D141A simulations (2 µs). Blue lines/bands: WT (solid line: average RMSD; shaded region: standard deviation); Orange lines/bands: D141A. (**G**) Bar graphs comparing how often HT-1 forms ionic interactions with key residues (D141, E33, D377 of BmrC and E297 of BmrD). WT shows a measurable interaction with D141. The mutant compensates for the missing interaction to some degree by forming more frequent ionic contacts with other residues (e.g., E33 of BmrC and D44 of BmrD), reflecting a shift in electrostatic contacts once D141 is removed. (**H**) Depicts ionic interactions of HT-2 with select negatively charged residues (e.g., D377, D45, D44 etc.). In WT, the distribution is typically anchored by D141’s network. In the mutant, D141 is no longer involved, and HT-2 shifts its interactions to other nearby residues (eg., E51).

**Fig. S10.**
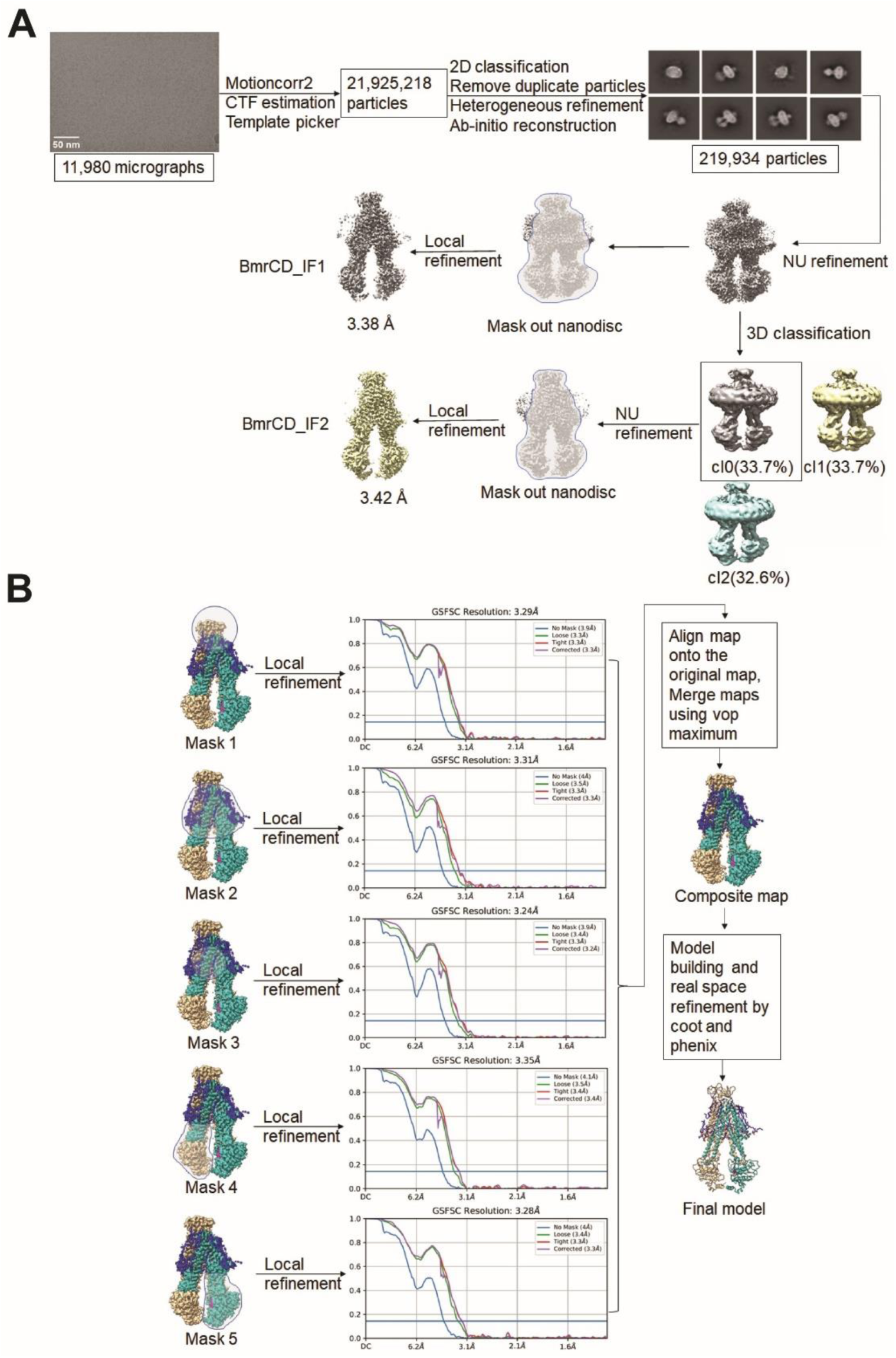
Cryo-EM data processing for BmrCD_IF1 and IF2. (**A**) Cryo-EM data processing workflow for BmrCD_IF1 and IF2. Data were processed by cryoSPARC. Representative micrographs, 2D, and 3D images are shown above. (**B**) Local refinement strategy for BmrCD_IF1. Masks is shown in transparent.

**Fig. S11.**
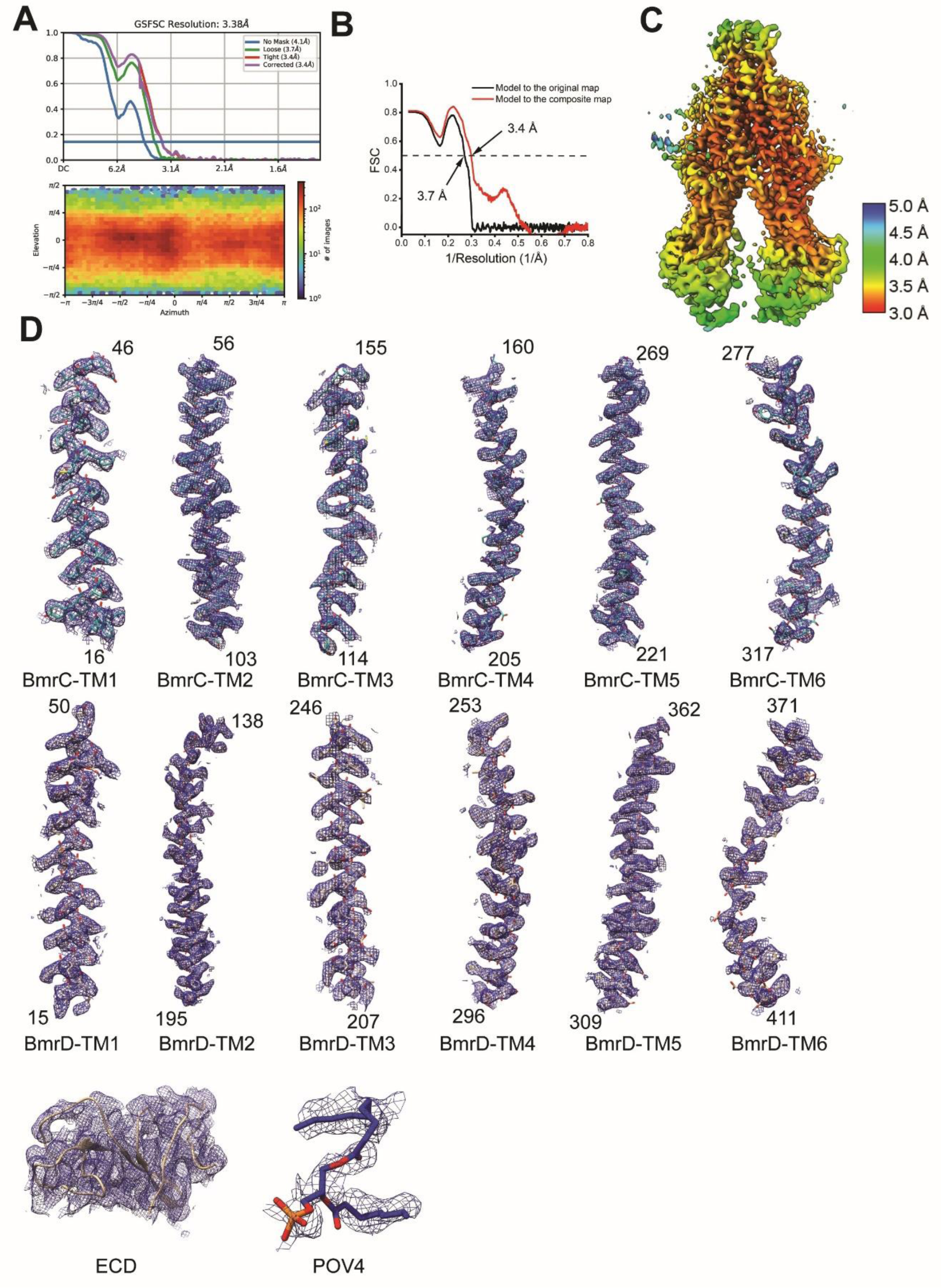
Cryo-EM data processing results for BmrCD_IF1. (**A**) Fourier shell correlation (FSC) curves (upper panel) and angular distribution (lower panel) of original map of BmrCD_IF1 after local refinement in cryoSPARC. (**B**) Fourier shell correlation (FSC) curves of the refined model versus the original map of BmrCD_IF1 after Non-uniform refinement in cryoSPARC (black) and the composite map (red) are plotted. (**C**) Local resolution maps for original maps of BmrCD_IF1 after local refinement in cryoSPARC. (**D**) Representative cryo-EM densities: transmembrane helices of BmrC and BmrD (residue numbers are indicated), extracellular domain (ECD), and lipid molecule POV4.

**Fig. S12.**
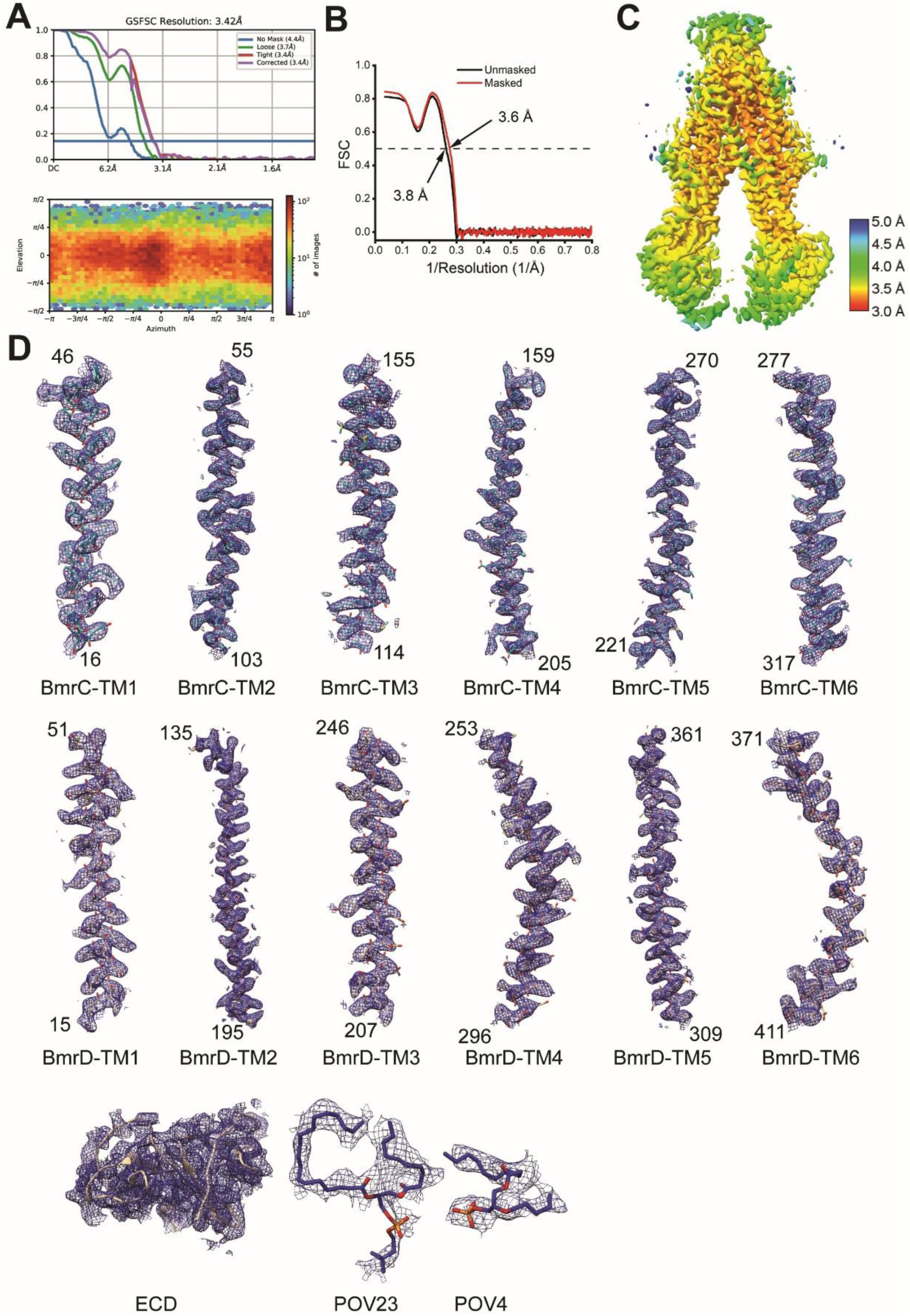
Cryo-EM data processing results for BmrCD_IF2. (**A**) Fourier shell correlation (FSC) curves (upper panel) and angular distribution (lower panel) of original map of BmrCD_IF2 after local refinement in cryoSPARC. (**B**) Fourier shell correlation (FSC) curves of the refined model versus the original map of BmrCD_IF2 after Non-uniform refinement in cryoSPARC are plotted. (**C**) Local resolution maps for original maps of BmrCD_IF2 after local refinement in cryoSPARC. (**D**) Representative cryo-EM densities: transmembrane helices of BmrC and BmrD (residue numbers are indicated), extracellular domain (ECD), and lipid molecules POV4 and POV23.

**Fig. S13.**
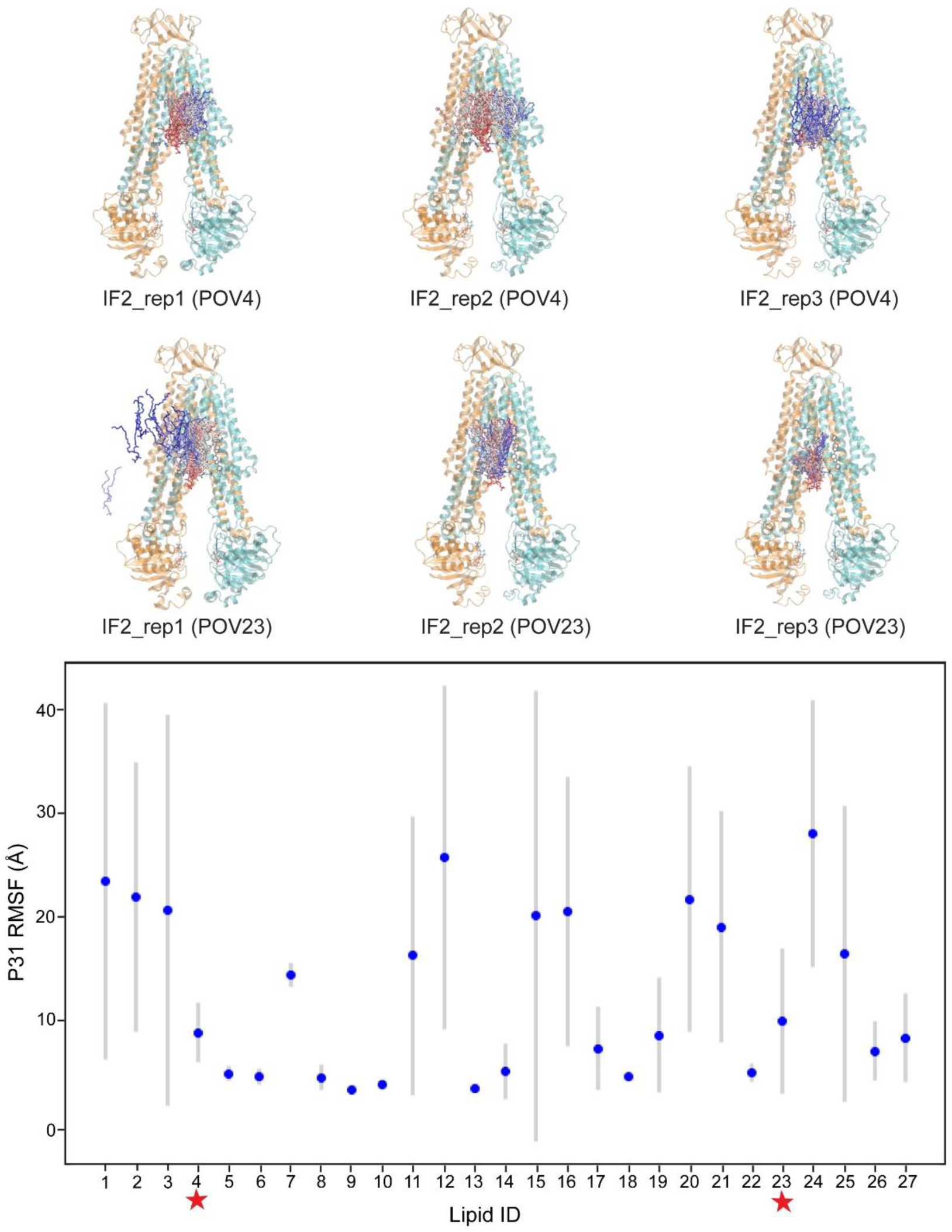
Flexibility of bound lipid heads. Atomic fluctuations plotted as average and standard deviations of the phosphorous atoms for all bound lipid densities over 3 replicas of 4 μs of molecular time each (aggregated 12 μs). The lipids (POV4 and POV23, denoted by star in the plot) that were found bound at the TM4/TM6 interface are not the most stable. Snapshots of such lipids from each replica for the lipid-bound simulation are shown as sticks and colored by timestep from red to blue. Both lipids progressively leave their site. POV4 leaves its site in IF2_rep2 and POV23 leaves its site in IF2_rep1.

**Fig. S14.**
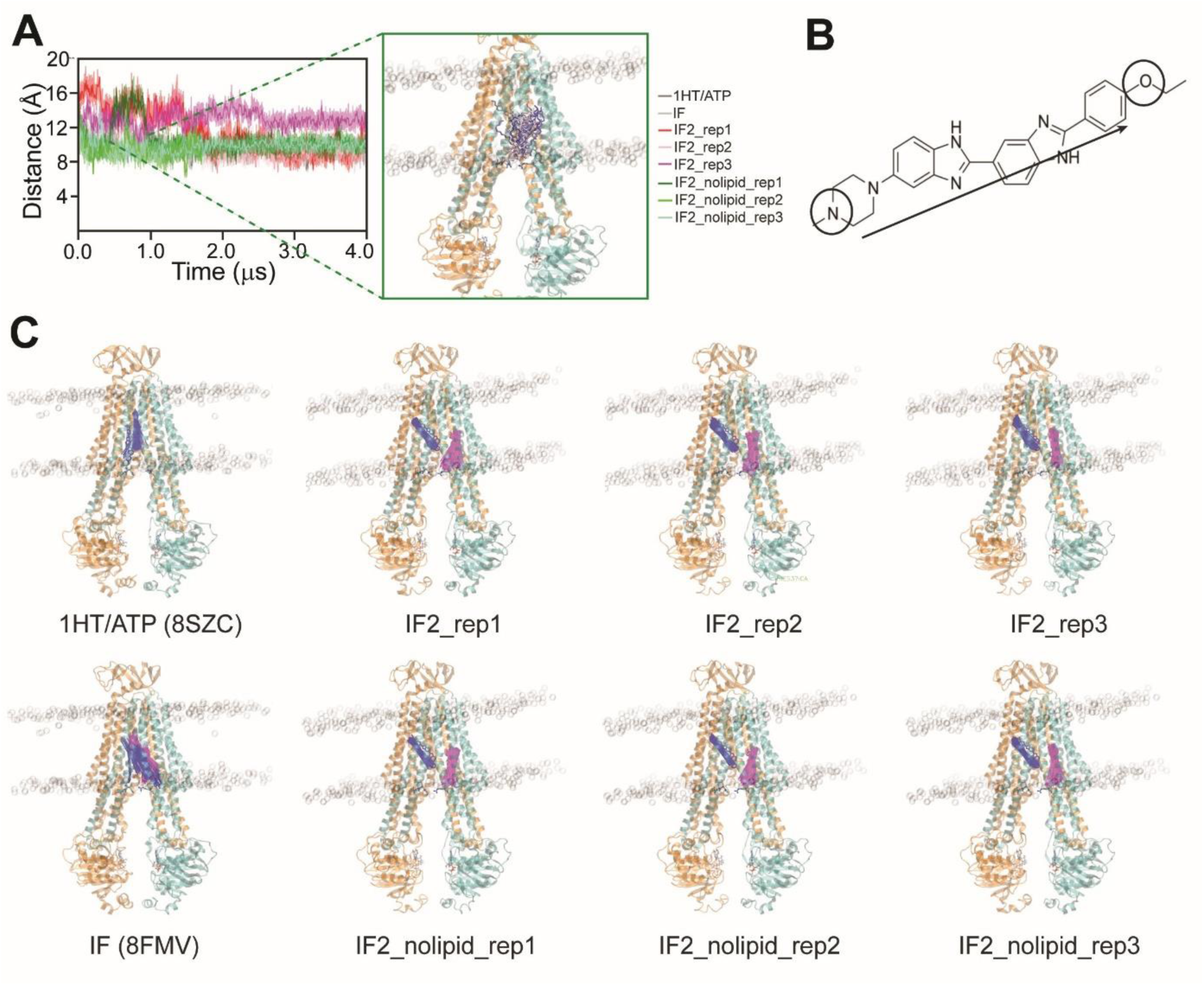
Snapshots for the MD simulations for Fig. 4. (**A**) Time series of the distance between the two RH-latch residues. In one of the BmrCD_IF2 simulations, a lipid molecule interacts at the TM4/TM6 interface after 400 ns (highlighted in inset). (**B**) and (**C**) HT vector connects the piperazine to the ethoxy group (**B**). Snapshots of HT molecules (HT-1 is colored by blue and HT-2 is magenta) during the simulations are shown as vectors for all simulated conditions, including all replicas (**C**).

**Fig. S15.**
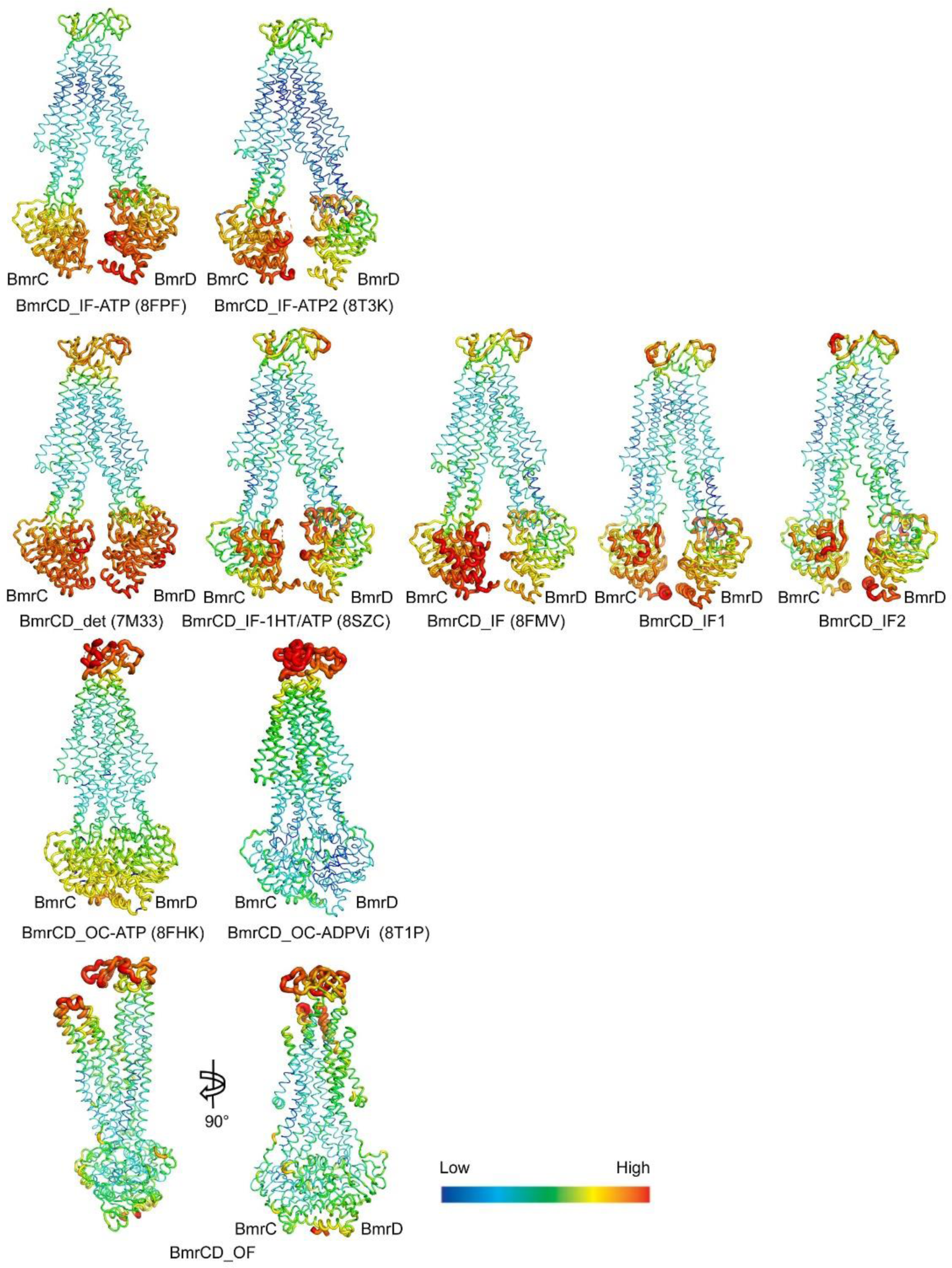
B-factor distribution analysis for different BmrCD conformations. Higher B-factor values are represented by red and thicker regions, while lower B factor values are represeted by blue and thinner regions. PDB IDs are listed in parentheses(*8, 21*). BmrCD_IF, BmrCD_IF1, BmrCD_IF2, and BmrCD_OF all have two Hoechst molecules bound. The comparesion between Hoechst-free and Hoechst-bound IF conformations suggests that Hoechst may increase the dyanmics of the ECD. The NBD of BmrC is more flexible than that of BmrD.

**Fig. S16.**
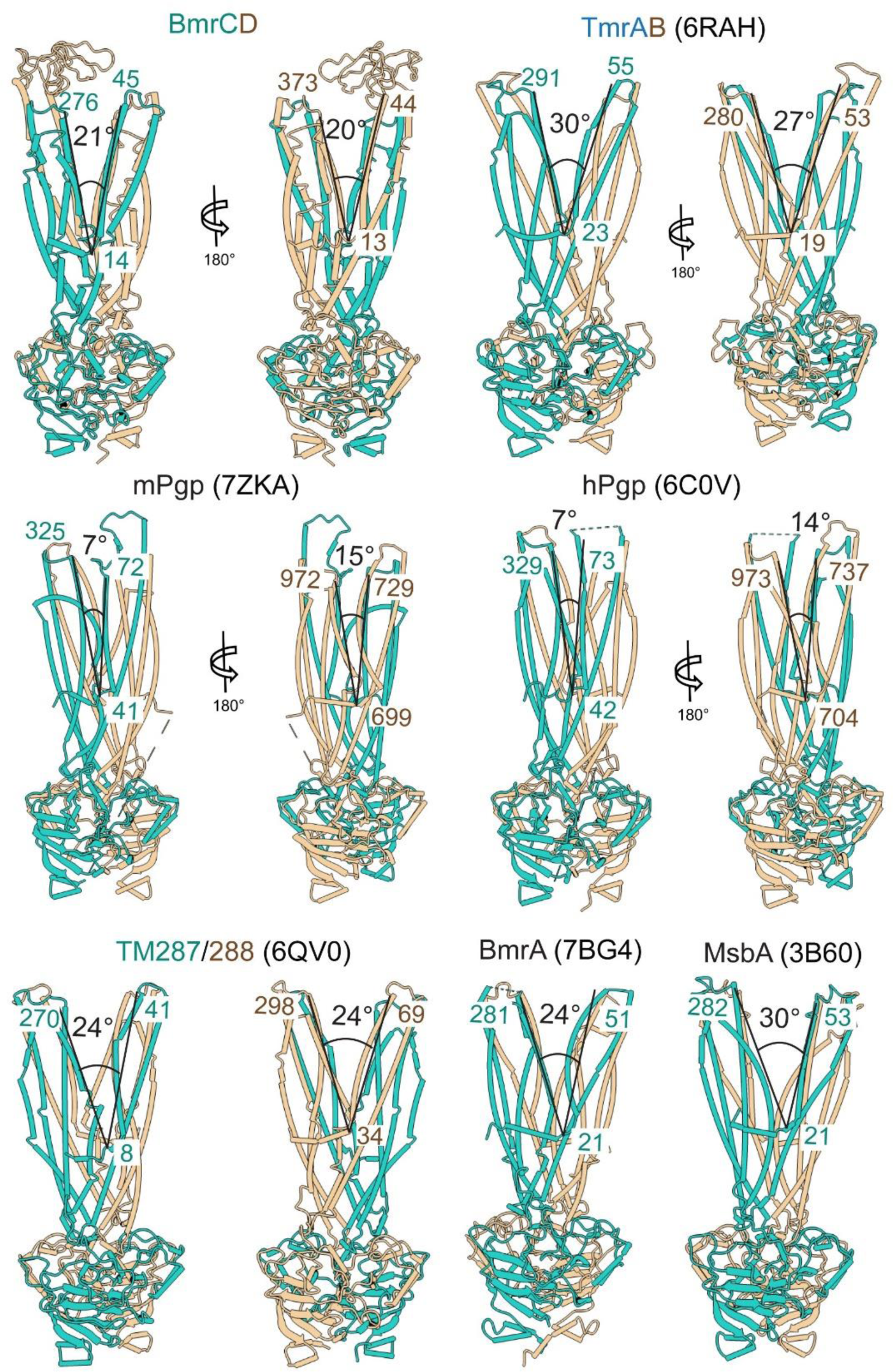
The opening angle at the extracellular side for the OF conformation of some representative ABC transporters. PDB IDs(*7, 10, 16, 63*) are listed in parentheses. The IDs of residues that used for measurement are labeled. BmrC, TmrA, N-terminals of mPgp and hPgp, TM287, and one subunit of BmrA and MsbA are colored in light sea green, while the others are colored in tan.

**Fig. S17.**
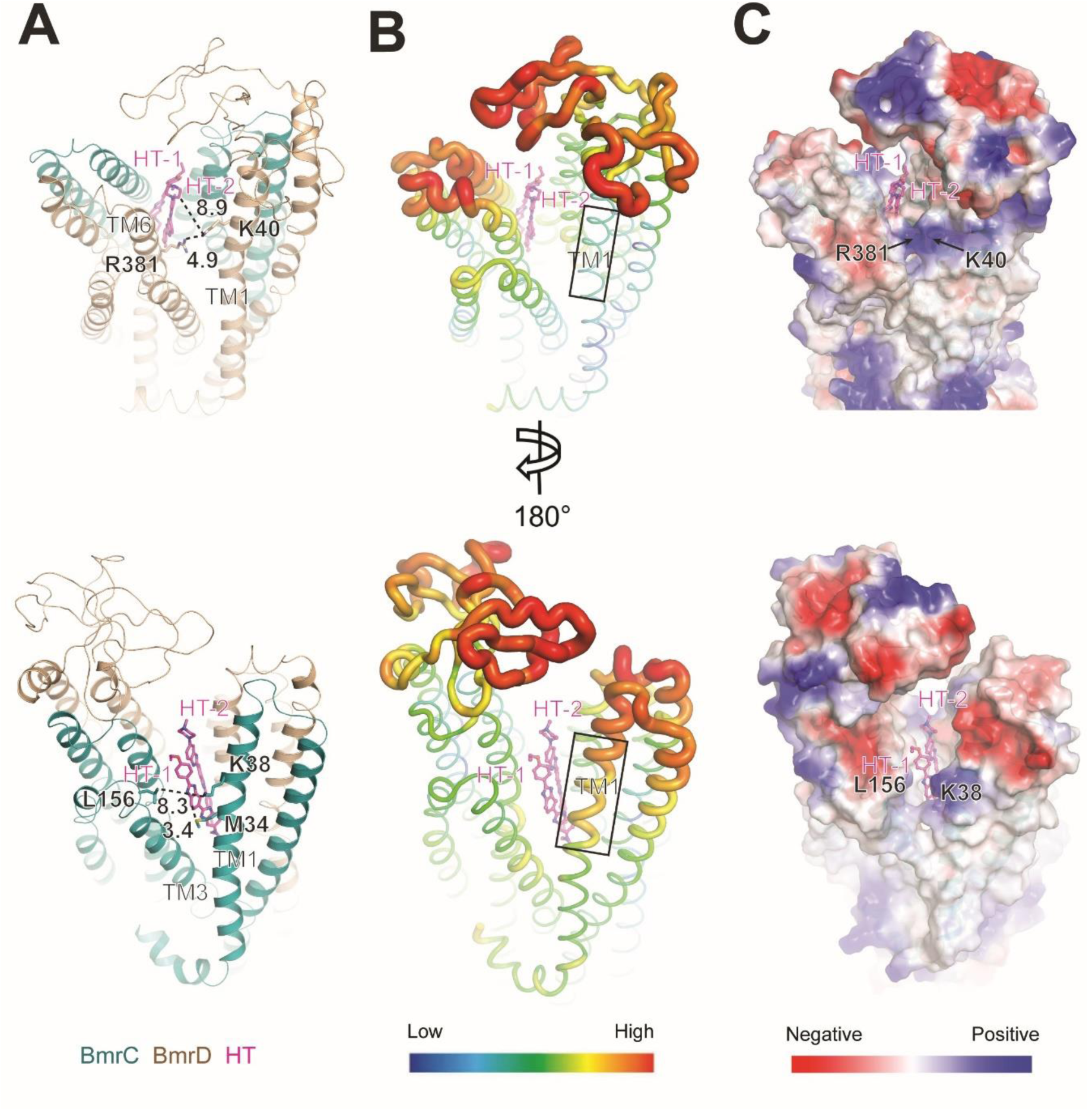
Analysis of the extracellular side of BmrCD_OF. (**A**), (**B**) and (**C**) are cartoon representation, B-factor distribution and electrostatics distribution of BmrCD_OF extracellular side. The upper and lower panels corresponidng to sideviews of BmrD and BmrC, respectively. Key residues are shown in stick representation. Distances are indicated in Å, and black rectangle highlights the comparsion region of TM1.

**Table S1.** Cryo-EM data collection, refinement, and validation statistics.

| **Data collection and processing** |  |  |  |
| --- | --- | --- | --- |
| Conformational state | BmrCD_OF | BmrCD_IF1 | BmrCD_IF2 |
| PDB ID | 9CUS | 9CUP | 9CUR |
| EMDB ID | EMD-45940 | EMD-45938 | EMD-45939 |
| Symmetry imposed | C1 |  |  |
| Microscope | Titan Krios (FEI) |  |  |
| Detector | Gatan K3 |  |  |
| Nominal magnification | 130,000 x |  |  |
| Voltage (kV) | 300 |  |  |
| Electron exposure (e/Å<sup>2</sup>) | 56 |  |  |
| Defocus range (μm) | -0.9 to -2.0 |  |  |
| Pixel size (Å) | 0.647 |  |  |
| Number of Micrographs | 11,980 |  |  |
| Particles images | 6,565,035 | 21,925,218 |  |
| Final particles images | 27,802 | 219,934 | 74,260 |
| Map resolution (Å) (FSC threshold=0.143) | 3.1 | 3.4 | 3.4 |
| **Refinement** |  |  |  |
| Model resolution (Å) (original map, FSC threshold=0.5) | 3.3 | 3.7 | 3.6 |
| Model resolution (Å) (composite map, FSC threshold=0.5) | 3.3 | 3.4 | N.A. |
| B-factor used for map sharpening (Å<sup>2</sup>) | -51.8 | -75.3 | -85.2 |
| **Model composition** |  |  |  |
| Non-hydrogen atoms | 10,754 | 10,948 | 10,831 |
| Protein residues | 1,244 | 1,246 | 1,225 |
| ATP, AOV, Hoechst, lipids, Mg<sup>2+</sup> | 1, 1, 2, 20, 2 | 2, 0, 2, 26, 1 | 2, 0, 2, 27, 1 |
| **Mean B factors (Å<sup>2</sup>)** |  |  |  |
| Protein | 85.88 | 66.56 | 66.52 |
| Ligands | 116.60 | 64.03 | 72.59 |
| **R.m.s. deviations** |  |  |  |
| Bond lengths (Å) | 0.003 | 0.009 | 0.011 |
| Bond angles (°) | 0.473 | 0.717 | 0.903 |
| Molprobity score | 1.45 | 1.74 | 1.66 |
| Clash score | 3.66 | 5.38 | 6.31 |
| Poor rotamers (%) | 0.00 | 0.28 | 0.00 |
| **Ramachandran plot** |  |  |  |
| Favored (%) | 95.81 | 92.99 | 95.47 |
| Allowed (%) | 4.19 | 7.01 | 4.53 |
| Disallowed (%) | 0.00 | 0.00 | 0.00 |

**Table S2.** Distance (Å) between COM of HT-1/HT-2 and residue D141 (or A141 in the mutant) and average RMSD of HT-1/HT-2.

| Ligand | System | Avg. Dist. to 141 (Å) | Std. Dev (Å) | Range (Å) |
| --- | --- | --- | --- | --- |
| HT-1 | WT | 4.0 | ± 1.7 | 1.5–9.9 |
| HT-1 | D141A | 7.5 | ± 2.5 | 2.0–14.4 |
| HT-2 | WT | 8.4 | ± 2.7 | 2.3–16.8 |
| HT-2 | D141A | 12.0 | ± 1.8 | 2.5–18.8 |
| Ligand | System | Avg. RMSD (Å) | Std. Dev (Å) | Range (Å) |
| HT-1 | WT | 3.8 | ± 1.0 | 0–7.6 |
| HT-1 | D141A | 6.1 | ± 3.1 | 0–12.3 |
| HT-2 | WT | 3.9 | ± 1.6 | 0–7.5 |
| HT-2 | D141A | 4.6 | ± 1.3 | 0–10.2 |

**Table S3.** Simulation setups for BmrCD_IF2.

| Name | Nucleotide(s) | Mg<sup>2+</sup> | Molecular Time (μs) | Substrate | Bound lipids | Composition | System Size |
| --- | --- | --- | --- | --- | --- | --- | --- |
| BmrCD_1HT/ATP (8SZC) | 2 ATP | 1 (consensus site) | 4.0 | 1 HT | no | 10% PA<br>90% PC (like experiment) | 342,169 |
| BmrCD_IF (8FMV) | 2 ATP | 1 (consensus site) | 4.0 | 2 HT | no | 10% PA<br>90% PC (like experiment) | 342,231 |
| BmrCD_IF2_rep1 (replica 1) | 2 ATP | 1 (consensus site) | 4.0 | 2 HT | 27 molecules | 10% PA<br>90% PC (like experiment) | 368,005 |
| BmrCD_IF2_rep2 (replica 2) | 2 ATP | 1 (consensus site) | 4.0 | 2 HT | 27 molecules | 10% PA<br>90% PC (like experiment) | 368,005 |
| BmrCD_IF2_rep3 (replica 3) | 2 ATP | 1 (consensus site) | 4.0 | 2 HT | 27 molecules | 10% PA<br>90% PC (like experiment) | 368,005 |
| BmrCD_IF2_nolipid_rep1 (replica 1) | 2 ATP | 1 (consensus site) | 4.0 | 2 HT | no (removed) | 10% PA<br>90% PC (like experiment) | 364,220 |
| BmrCD_IF2_nolipid_rep2 (replica 2) | 2 ATP | 1 (consensus site) | 4.0 | 2 HT | no (removed) | 10% PA<br>90% PC (like experiment) | 364,220 |
| BmrCD_IF2_nolipid_rep3 (replica 3) | 2 ATP | 1 (consensus site) | 4.0 | 2 HT | no (removed) | 10% PA<br>90% PC (like experiment) | 364,220 |

**Table S4.** Root mean square deviation (r.m.s.d.) of overall structure (white, above diagonal) and TMD (grey, below diagonal).

|  | BmrCD_OF | TmrAB_OF | mPGP_OF | hPgp_OF | TM287/288_OF | MsbA_OF | BmrA_OF |
| --- | --- | --- | --- | --- | --- | --- | --- |
| BmrCD_OF | 0.00 | 1.6 (904 Cα) | 1.6 (758 Cα) | 2.4 (838 Cα) | 1.5 (879 Cα) | 1.3 (794 Cα) | 2.0 (844 Cα) |
| TmrAB_OF | 2.1 (470 Cα) | 0.00 | 1.6 (758 Cα) | 2.3 (801 Cα) | 1.5 (997 Cα) | 1.5 (845 Cα) | 2.2 (854 Cα) |
| mPgp_OF | 3.6 (439 Cα) | 3.5 (440 Cα) | 0.00 | 1.5 (1031 Cα) | 1.6 (783 Cα) | 1.8 (773 Cα) | 1.8 (769 Cα) |
| hPgp_OF | 3.7(463 Cα) | 4.1 (457 Cα) | 1.4 (564 Cα) | 0.00 | 2.2 (822 Cα) | 2.2 (769 Cα) | 2.2 (777 Cα) |
| TM287/288_OF | 2.7 (533 Cα) | 1.7 (558 Cα) | 3.1 (423 Cα) | 3.4 (409 Cα) | 0.00 | 1.4 (857 Cα) | 1.8 (831 Cα) |
| MsbA_OF | 3.9 (530 Cα) | 2.4 (497 Cα) | 5.0 (454 Cα) | 5.0 (428 Cα) | 2.5 (499 Cα) | 0.00 | 1.6 (865 Cα) |
| BmrA_OF | 2.7 (440 Cα) | 3.6 (545 Cα) | 3.9 (393 Cα) | 4.5 (441 Cα) | 3.0 (499 Cα) | 2.1 (438 Cα) | 0.00 |

Superposition was performed at the residue level by Pymol and r.m.s.d. was calculated at the Cα level in unit of Å. The PDB IDs are: TmrAB\_OF, 6RAH; mPgp\_OF, 7ZKA; hPgp\_OF, 6C0V; TM287/288\_OF, 6QV0; MsbA\_OF, 3B60; BmrA\_OF, 7OW8.

**Data S1. (separate file)**

Initial coordinates, simulation input files and the final output files for BmrCD_OF.

**Data S2. (separate file)**

Initial coordinates, simulation input files and the final output files for BmrCD IF structures.

